# A mechanical model of multicellular remodelling in epithelial monolayers

**DOI:** 10.64898/2026.09.09.750324

**Authors:** James M. Osborne, Reuben Van Ammers, Yohan Davit, David J. Gavaghan, Helen M. Byrne

## Abstract

Epithelial monolayers are the foundation of many mammalian organs and have a significant impact on numerous biological processes, including the secretion of cytokines, absorption of waste products and barrier function. During organ development epithelia are subject to many external forces due to the growth of the surrounding tissues. They exhibit complex responses to such forces, with deformation occurring on short timescales (seconds), and relaxation and remodelling occurring on longer timescales (minutes or hours).

Existing mathematical and computational models do not typically account for subcellular remodelling, assuming instead that the response to an applied force or strain acts on a single timescale. In this paper we extend an off–lattice, cell–centre modelling framework to account for subcellular and tissue remodelling by introducing a Dynamic Reference Frame (DRF). The DRF provides a discrete, cell–based analogue of morpho–elasticity: it decouples the evolution of each cell’s mechanical reference configuration from its instantaneous deformation, providing a phenomenological description of subcellular remodelling processes — cytoskeletal reorganisation, myosin turnover and adhesion bond remodelling — at the cellular scale. Using this extended multicellular model, we reproduce multiscale responses to external mechanical forces, including creep and stress relaxation experiments, and demonstrate that the history of applied deformation influences tissue recovery upon release. Additionally we identify and quantify where and how these multiscale responses occur. The resulting framework allows for more detailed descriptions and analyses of the development and function of biological tissues.

## 1. Introduction

Epithelial tissues form the building blocks for mammalian organs. Their primary functions are to protect vital organs (through barrier function), secrete hormones and enzymes, and to absorb harmful substances. Arguably, the simplest of these tissues are the epithelial monolayers that line many organs [1], including: the intestines [2]; the respiratory tract [3]; blood vessels [4]; and the kidneys [5]. As well as being important during healthy development, epithelial tissues are often initiating sites of cancer, partly due to their high rates of cell division and direct exposure to environmental mutagens [1].

Experimental studies of epithelial monolayers have demonstrated that their response to deformation can span multiple timescales, from seconds to minutes (or longer), even when cell proliferation and rearrangement are negligible [6, 7]. To better understand this effect, creep and stress relaxation experiments have been performed [6] (see Figure 1). In a creep experiment, a constant force is applied and strain is recorded, for low (blue) and high (red) loads (Figure 1 (a)). In a stress relaxation experiment, the tissue is stretched to a fixed strain and the resulting stress recorded (Figure 1 (b), note, the plotted quantity is the excess stress, *σ − σ_∞_*, where *σ_∞_* is the long–time stress) with traces corresponding to low (blue) and high (red) applied strain rates. In both cases, for certain conditions (creep with 3.0 kPa, and stress relaxation at a rate of 0.75 mms*^−^*^1^), the tissue response spans multiple timescales: a fast elastic response on the order of seconds and a slower relaxation or creep response on the order of minutes [6].

**Figure 1:**
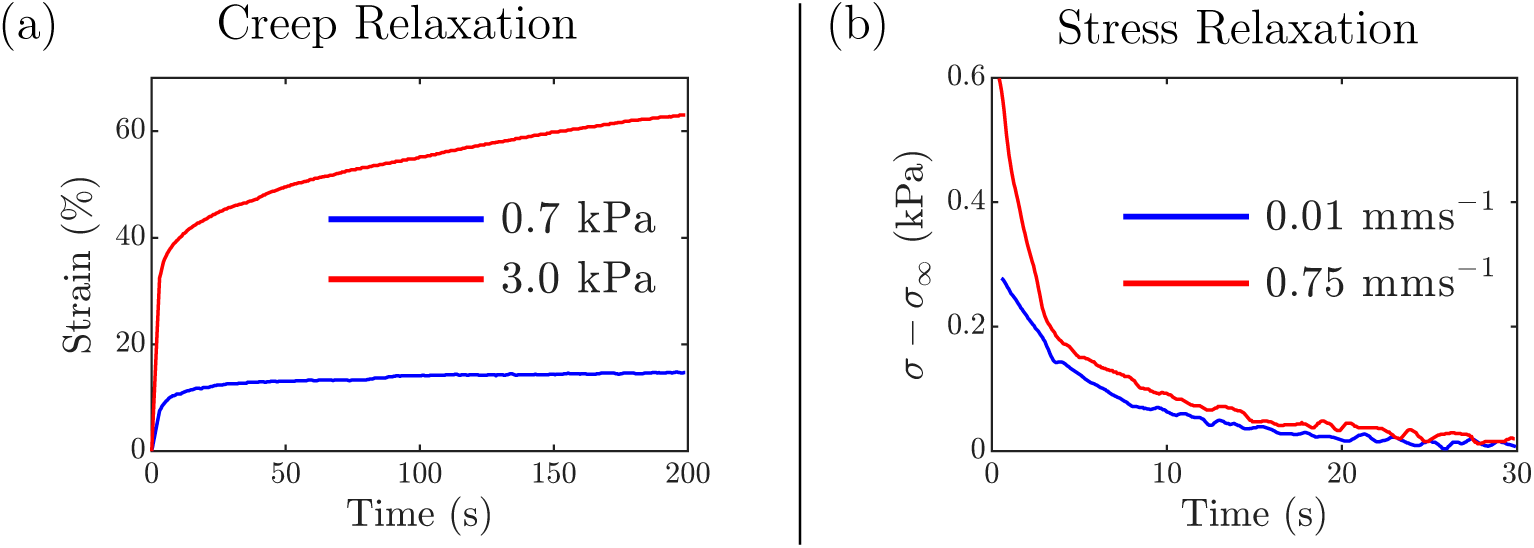
Experimental evidence for multiscale responses to applied deformations. Summary of experiments from Harris et al. [6]. (a) Strain over time of a biological monolayer in response to a low (blue) and high (red) external stress in a creep relaxation experiment. (b) Stress (*σ − σ_∞_*) over time following low, (blue) and high, (red), strain–rate loading respectively for a stress relaxation experiment. Figures generated from the experimental data presented in Harris et al. [6].

These experiments are performed on timescales that are too short for cellular–level reorganisations or growth to occur. As a result the cause of the multiple timescale response is not completely clear. Cells are deformable and their mechanical properties (and, therefore, those of the tissues in which they reside) are strongly dependent on: the integrity of each cell’s actin cytoskeleton; the level of myosin within each cell; and the inter–cellular adhesion between neighbouring cells [6]. Our working hypothesis is that remodelling and turnover of specific subcellular components — in particular the actin cytoskeleton, myosin activity, and cell–cell adhesion bonds — in response to applied stress or strain alters the mechanical properties of individual cells. These changes subsequently give rise to deformation responses that act on longer timescales. In this paper we introduce a modelling framework that incorporates such effects at the multicellular scale, without requiring explicit representation of subcellular mechanics. To place our approach in context, we now review existing mathematical models that have been used to describe the interplay between tissue mechanics and deformation.

A standard approach to modelling tissue mechanics involves using continuum models based on partial differential equations where the tissue may be represented as an elastic or viscoelastic material. The simplest viscoelastic model, based on a superposition of linear springs and dashpots (one spring in parallel with a spring and dashpot system in series), is known as the standard linear solid (SLS) model [8]. Here, creep and stress relaxation responses decay exponentially over time. Therefore, the deformation evolves with a single characteristic timescale. More complex viscoelastic models, such as those comprising multiple Maxwell elements (a spring and dashpot in series) in series, can in principle produce responses on multiple timescales by construction; however, such models are largely phenomenological, with parameters that are challenging to relate directly to subcellular mechanisms [9].

More detailed morpho–elastic models decouple elastic deformation from deformation due to tissue growth, allowing the mechanical properties of the material to change over time and capturing multiple timescales in the tissue response [10]. Morpho–elasticity has been used to understand, among others: tissue deformation during wound healing [11]; tissue formation by stem cells [12]; ventricular hypertrophy [13]; and buckling of airways [14]; see also the reviews by Goriely et al. [15] and Ambrosi et al. [16] for a comprehensive treatment of the mathematical foundations and biological applications of continuum modelling of tissue growth and development. In these systems, the multiple timescales arise from differences between the timescales of tissue growth (and/or cell turnover) and of mechanical deformation of the evolving tissue.

Multicellular or cell–based approaches are also used to model tissue mechanics, treating cells (and/or their subcellular components) as discrete, mechanically interacting entities. As such, they provide a natural framework for studying the regulation of cell–level processes in tissue and organ development, allowing a one–to–one correspondence between cells in the model and the tissue [17]. This enables tissue heterogeneity (due, for example, to mutations in individual or small populations of cells, or to heterogeneity in the spatial distribution of signalling factors) to be incorporated by specifying appropriate cellular interactions. This allows heterogeneity in processes such as secretion and absorption of enzymes and morphogens to be captured [17]. Existing cell–based models vary in complexity; here we restrict our attention to off–lattice models, which are based on mechanical interactions [17] (see Table 1 for a summary of these models and their properties relevant to representing remodelling).

**Table 1:** Discrete cell–based model summary. Discrete (cell–based) tissue models representing tissue deformation. Properties relevant to representing subcellular remodelling are highlighted. DoF = Degress of Freedom, Tissue–Scale = 1000s of cells in 2D and or 3D.

| Model | Cell Representation | Relative DoF / Cost | Can Represent Subcellular Remodelling? | Scalable to Tissue-Scale? | Refs |
| --- | --- | --- | --- | --- | --- |
| Overlapping Spheres (OS) | Point at cell centre | Low | No — fixed rest length/radius | Yes | [17, 18] |
| Voronoi Tessellation (VT) | Point at cell centre; polygon from Voronoi region | Low-moderate | No — fixed rest length between centres | Yes | [17] |
| Vertex Dynamics (VD) | Polygon with moving vertices | Moderate-high | No — fixed target perimeter/area | Yes | [19] |
| Active Vertex Model (AVM) | Polygon with moving vertices | Moderate-high | No — solid/fluid transition set by a fixed shape index, not dynamic | Yes | [25, 26] |
| Mechanosensitive Vertex Dynamics | Polygon with moving vertices | Moderate-high | Partial — dynamic parameters depend on stress via prescribed functional relations | Yes | [27] |
| Edge Based (EB) | Collection of moving edges to create polygon cells | Moderate | No — fixed interaction forces | Yes | [23] |
| Subcellular Element (SE) | Multiple points per cell (internal and intercellular interactions) | High | Yes — in principle, via internal reorganisation | No (<10 cells) | [21] |
| Finite Element (FE) | Interconnected triangular/tetrahedral elements | High | No — standard elastic constitutive law | Limited (mesh cost) | [22] |
| Hyperelastic Chain | Discrete 1D chain, hyperelastic springs | Low | No — fixed hyperelastic constitutive law | 1D only | [28] |
| Biphasic Chain | Discrete 1D chain, active/passive spring phases | Low | Yes — via phase-dependent thresholds (many coupled parameters) | Yes (2D/3D possible, in principle) | [29] |

A popular class of off–lattice models are cell centre based models, wherein cell locations are represented by points located at the centers of each cell. Forces due to external factors, and cell–cell interactions, are assumed to act at these cell centres. Depending on their connectivity, cells can be represented as (a set of) Overlapping Spheres (known as the OS model, or as the Point Force model [18]) or as (a set of) polygonal cells using the Voronoi Tessellation (or VT) model [17]. Vertex Dynamics (or VD) models, represent cells as polygons whose vertices move [19]. They have increased computational cost but offer more flexibility in describing systems than cell centre based models, because they have more degrees of freedom [20]. Other cell–based models represent cells at higher levels of detail (and even greater computational cost). The Subcellular Element (SE) model, for example, uses multiple points to represent a cell’s internal elements and distinguishes between interactions involving points within a cell and those involving multiple cells or components [21]. The Finite Element (FE) model represents cells using interconnected triangular (or tetrahedral) elements which can deform [22]. Finally the Edge Based (EB) model represents cells as polygons, much like the VD model. Here, however, the cells are represented as a collection of edges [23]. In their standard formulation, when cell proliferation and rearrangement are neglected and elastic interaction laws are used at the smallest spatial scale, the response of the above multicellular models, is dominated by a single timescale [24]. The SE model is a potential exception: by resolving multiple interacting components within each cell, it could in principle capture responses on multiple timescales through subcellular reorganisation [21]. However, the associated computational cost restricts its application to small systems (less than 10 cells), making it impractical for the tissue–scale simulations considered here.

Another approach to adding higher order behaviour to centre based multicellular models is to edit the form of the interaction force between cells to more accurately capture observed behaviour. Barry et al. developed a non–linear partial differential equation model by upscaling a one dimensional model of a chain of deformable cells with hyperelastic interactions [28]. However, this model is not directly extensible to higher spatial dimensions due to the form of the cellular interaction. In separate work, Germano et al. used a biphasic spring model to account for cell–cell interactions and used their model to simulate plastic deformation of a chain of cells. As with the Maxwell–element models above, extension of this model to higher dimensions is possible, but requires the specification of multiple coupled parameters (spring stiffnesses, drag coefficients and remodelling thresholds for each active and passive phase) that are difficult to measure independently [29].

Arguably, VD models are the most popular framework for studying viscoelastic and morpho–elastic behaviour in multicellular tissues. In a series of papers, Jensen et al. have investigated the mechanical properties of the VD model and used it to reproduce experimentally observed stress distributions [30–32]. An extension of the VD model known as the Active Vertex Model (AVM) can exhibit solid–like (jammed) or fluid–like (unjammed) behaviour, depending on the mean cell shape index [25, 26]. A given tissue operates in one of these two regimes; the transition between them requires a change in the shape index parameter and does not emerge dynamically from the equations of motion. Another extension of the VD model accounts for mechanosensitive junction remodelling by allowing model parameters to depend on the stress distributions in the tissue through experimentally motivated functional relations. While promising, it is challenging to identify all possible model behaviours [27].

Non–linear viscoelastic models based on fractional derivatives have also been used to capture multiscale tissue rheology, treating the tissue as a one–dimensional continuum rather than a collection of individual cells [33, 34]. However, their inherently integro–differential, global memory–kernel formulation makes extension to higher dimensions difficult.

In spite of their strengths and previous accomplishments, the above approaches are either: (i) unable to capture the multiple timescales on which responses are seen in experiments, without cell rearrangement or proliferation; (ii) are limited to systems of a single (or two) cells; (iii) are limited to one–dimensional simplifications; (iv) contain functional relations or multiple parameters that are challenging to quantify; or (v) have a computational cost that limits their application to small systems or short timescales, making them impractical for tissue–scale simulations.

In this paper, we introduce a novel cellular level framework to explain the appearance of multiple timescales in stress and creep responses. In this framework cell–cell interaction forces do not change and proliferation and rearrangement are neglected. Instead, we incorporate a dynamic reference frame (DRF), allowing us to evolve the reference configuration for cell–cell interactions as a function of local stress and strain, into a multicellular model of a deforming tissue. We will illustrate the DRF using a Voronoi Tessellation model, although it can be applied to other off–lattice models, including those described above. Using the DRF, we can account for subcellular remodelling within individual cells, allowing them to respond to the stresses and strains that they experience. We also show how the properties of the DRF influence the timescales on which the tissue responds to stress and strain.

The remainder of the paper is structured as follows. First, in Section 2, we present the DRF and show how it can be used for multicellular simulations (the DRF model) of biological tissues. In Section 3, we present computational studies of the DRF model, which recapitulate creep and stress relaxation experiments. For each experiment, we show how varying the DRF model parameters influences the response timescales. We also show how the DRF enables the history of the deformation to influence how a tissue that has been subject to extension behaves upon release. Finally, in Section 4, we summarise our results, discuss their biological impact and present possible avenues for future work.

## 2. Methods

In this paper we focus on the deformation of a two–dimensional epithelial monolayer (of *N* cells) on timescales for which cell proliferation and rearrangement are negligible (these assumptions can be relaxed; for details, see Section 4). Here, we extend a cell–centre cell–based model based on Voronoi Tessellations to include a dynamic reference frame (DRF). The DRF allows a cell’s internal properties, specifically the rest length of its cell–cell interactions (which control cell size), to evolve in response to local applied stresses and displacements.

### 2.1. The Voronoi Tessellation model

In the Voronoi tessellation (VT) model, the tissue is represented as a set of physical cell centres *{****r****_i_}* where *i* = 1*, … , N* spans the monolayer. Cells are connected to their neighbours via a Delaunay Triangulation and cells are represented as polygons defined by the Voronoi region corresponding to each cell centre [17]. Newton’s Second Law, and the assumption that the system is over damped, are used to neglect inertial motion and balance internal linear spring–type forces and externally applied forces with viscous drag to derive the following equation of motion for each cell centre:

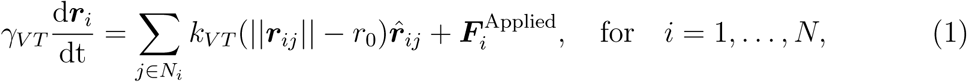

where *N_i_*is the set of neighbours of cell *i* (defined by the Delaunay Triangulation), and ***r****_ij_* = ***r****_i_ −* ***r****_j_*. We denote *||****r****_ij_||* to be the length of the “spring” between cell centres (calculated by the *l*_2_ norm) and *r*_0_ is the rest length of this “spring”. We also use hats to indicate unit vectors 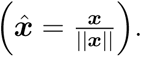. Any external force acting on cell *i* is denoted by 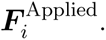. The drag coefficient for the cells *γ_VT_* controls the speed of cell movement and *k_VT_* is the spring constant defining the magnitude of interactions between cells. Equation (1) are solved subject to appropriate initial conditions to calculate the evolution of the physical cell centres {***r****_i_*} over time [17].

We now extend the VT model to include a dynamic reference frame (DRF) that allows the rest lengths of the cell–cell interactions to evolve in response to local stresses and strains.

### 2.2. The dynamic reference frame (DRF)

We extend the description of the tissue to be a set of physical cell centres {***r****_i_*} and corresponding reference cell centres {***ρ****_i_*}, which are associated with a set of static cell centres {***s****_i_*}, where *i* = 1*, … , N* spans the monolayer (see Figure 2 (a)). As in the VT model cells are connected to their neighbours via a Delaunay Triangulation based on the initial cell configuration (here a regular hexagonal lattice; see Section 2.3). Reference and static cells have the same connectivity as physical cells (see Figure 2 (b)). This connectivity defines the set of cell neighbours *N_i_* for each cell *i*, and is maintained throughout a simulation, so there is always a one–to–one correspondence between edges in each mesh as the physical and reference cells move.

**Figure 2:**
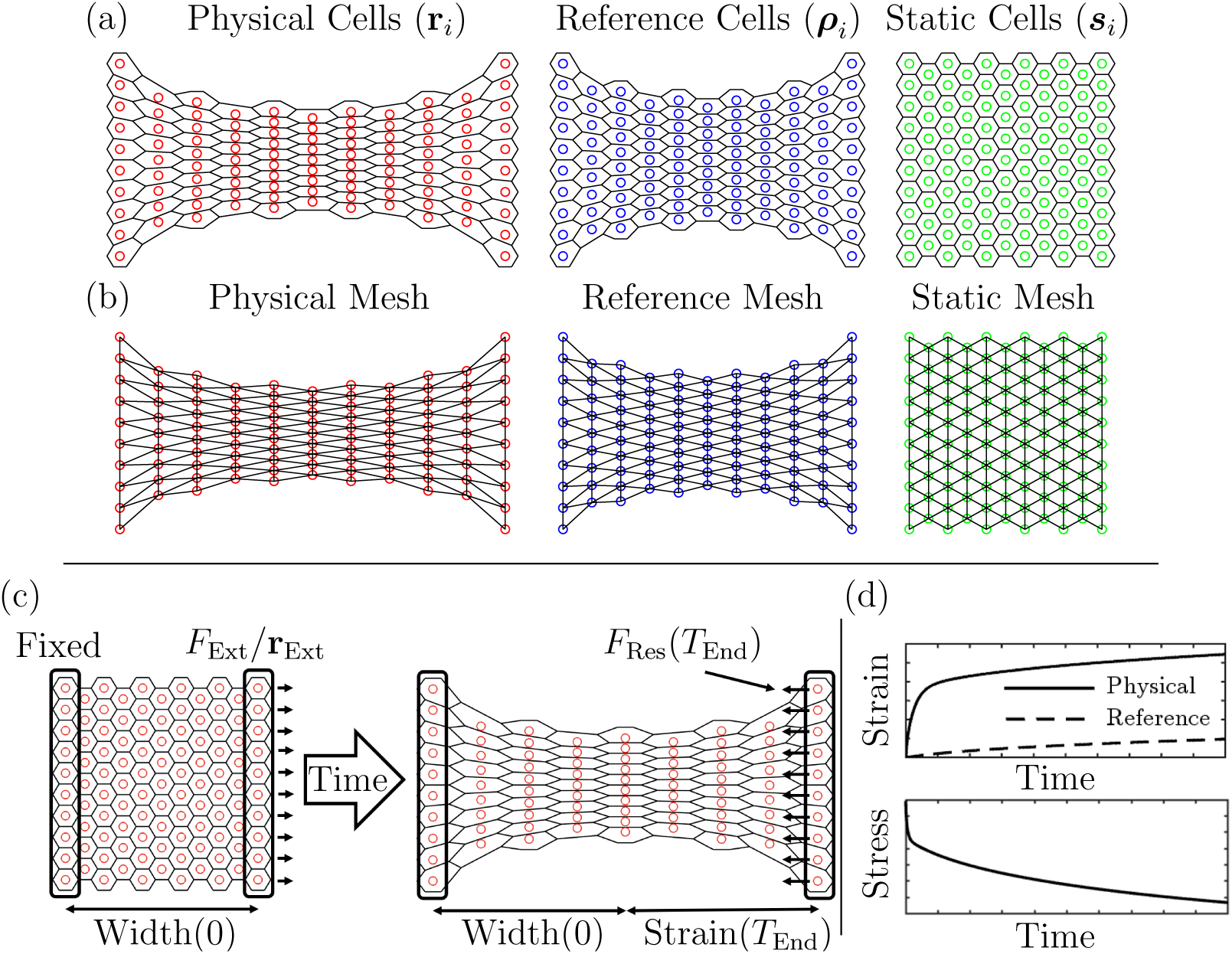
Illustration of reference frames in the model and simulation schematic. (a) and (b) are schematics of the three cell configurations used to describe the monolayer: physical, reference, and static. (a) shows the cell centre locations for each frame of reference and (b) shows the corresponding cell–cell connectivity or mesh. There is a one–to–one correspondence between cells in each frame of reference. (c) Experimental setup where a cell monolayer is held fixed on the left and a force (*F*_Ext_) or displacement (**r**_Ext_) is applied on the right edge. The initial configuration is shown on the left and the final deformed configuration is shown on the right. The strain (defined as the difference between the current tissue width and the initial tissue width normalised by the initial tissue width, indicated in (c) for both the physical and reference cells) and the resistive force *F*_Res_ (defined as the mean *x*–component of the net spring force acting on the right–boundary cells when held at their new positions) can be tracked over time to calculate the stress, and schematic examples of strain or stress over time are shown in (d).

As before, assuming overdamped dynamics and balancing internal spring–type forces (from neighbouring cells), externally applied forces, and viscous drag gives the following equation of motion for each physical cell:

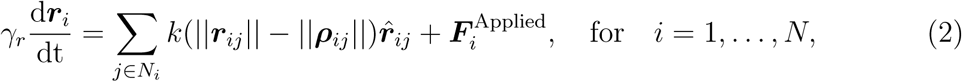

where *N_i_*is the set of neighbours of cell *i*, ***r****_ij_* = ***r****_i_ −* ***r****_j_* and ***ρ****_ij_* = ***ρ****_i_ −* ***ρ****_j_*. We denote by *||****r****_ij_||* and *||****ρ****_ij_||* the physical and reference lengths respectively. The external force acting on cell *i* is denoted by 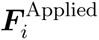 and depends on the experiment being simulated (see Section 2.3). The drag coefficient for physical cells *γ_r_* controls the speed of physical cell movement, while *k* is the spring constant for physical cells, here these are the same for all cells but could, in principle, vary. In the original VT model, Equation (1), *||****ρ****_ij_||* is replaced by a fixed reference length, *r*_0_. Instead, in our model, the position of reference cell centres ***ρ****_i_* = ***ρ****_i_*(*t*) can vary with time (and, therefore, *||****ρ****_ij_||* also varies). Using similar assumptions on movement, of reference cells, we introduce the following equation of motion for each reference cell:

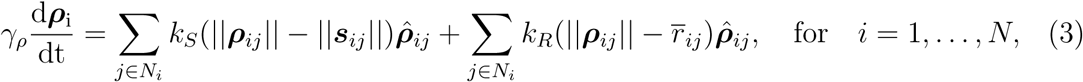

where ***s****_ij_* = ***s****_i_ −* ***s****_j_*, *||****s****_ij_||* denotes what we term the static length between cells *i* and *j* (which is constant in time), *γ_ρ_*denotes the drag coefficient for the reference cells, and *k_S_* and *k_R_* are spring constants relating to the static and remodelling components of the reference state, which control the influence of the remodelling and static contributions. Additionally, *r̄_ij_* is the average distance between physical cell centres *i* and *j* over the previous 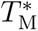 hours, used to capture the average tissue deformation, calculated as

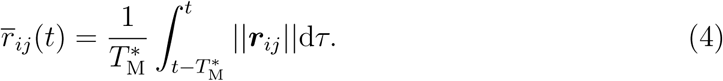

When defining *r̄_ij_*(*t*), we assume that the physical cells are stationary for *t ≤* 0 so that ***r****_ij_*(*t*) *≡ **r**_ij_*(0) for *t ≤* 0.

Equation (3) is a force balance for the reference frame. The first term on the right–hand side is a restoring force that drives the reference state towards the static configuration, while the remodelling force (second term) drives the reference state towards the mean physical configuration over the previous 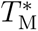 hours.

We use time–averaged distances *r̄_ij_* since we assume that the tissue remodels in response to prolonged deformation, rather than responding instantaneously. The specific functional form used to describe interactions between the physical and reference cells is chosen for illustrative purposes only. Other functional forms, where, for example, reference cell properties evolve in response to stress rather than strain, may also be considered. We note that Equations (2) and (3) reduce to the traditional VT model Equation (1) if *k_R_* = 0 or *k_S_ ≫ γ_r_* and *k_S_ ≫ k_R_*.

We assume that all physical and reference cells are initially at equilibrium in the static configuration, so that ***r****_i_*(0) = ***ρ****_i_*(0) = ***s****_i_* for all *i*. We denote the external forces and constraints that represent the experiment under consideration as boundary constraints. These boundary constraints are applied to the physical cells and are defined in Section 2.3 (see Figure 2 (c)).

Before proceeding further, it is convenient to non–dimensionalise the model variables as follows:

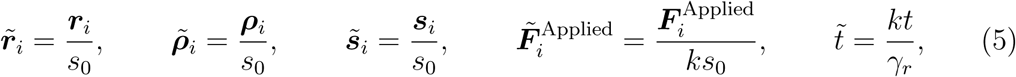

where *s*_0_ is the common static length *||**s**_ij_||*, which is constant in time, and the same for all connected static cell centres due to the regular hexagonal packing of the initial configuration (Section 2.3 and Figure 2 (c) right).

If we substitute from Equation (5) into Equations (2) and (3), and drop the tildes for notational simplicity, then we have:

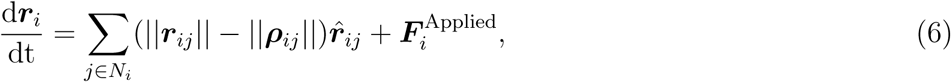

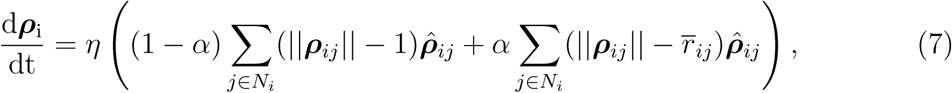

where

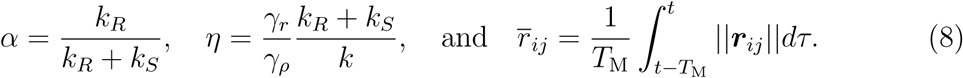

The dimensionless parameter *α* determines the relative strength of the static and active forces acting on the reference state (*α* = 0 corresponds to a static reference frame), the dimensionless parameter *η* determines the rate at which the reference state remodels compared to the rate at which the physical state evolves, and *T*_M_ is the dimensionless time over which the reference state responds to the physical state. (Note that where needed we denote dimensional variables with an *∗*; for example, 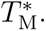.) We define *α* to be the remodelling ratio, *η* to be the remodelling rate and *T*_M_ to be the remodelling memory (initially we fix *T*_M_ = 0, however later in this study we relax this assumption).

### 2.3. Tissue setup, initial conditions, and boundary constraints

In all simulations we are working in two–dimensional Cartesian coordinates (*x, y*) with *x* pointing in the direction of the applied force or deformation. All experiments are performed on a hexagonal grid of 105 cells (11 columns, each comprising 9 or 10 cells, chosen so that the tissue is symmetric with respect to both the *x* and *y* axes) which are initially arranged in a regular configuration (Figure 2 (c), initial conditions). The tissue size was chosen to balance computational load with simulation fidelity, and we note that the qualitative results presented here remain unchanged for tissues with a larger number of cells. Cell connectivity is determined from the Delaunay Triangulation of the initial distributions of the cell centres and is maintained throughout each simulation (Figure 2 (b)). Cells on the left and right of the tissue are termed boundary cells (see Figure 2 (c)) and are subject to boundary constraints. Physical cells on the left of the tissue are held fixed and a force, *F*_Ext_, or displacement **r**_Ext_(*t*), is applied to physical cells on the right (Figure 2 (c), see Sections 3.1.1 and 3.2.1 for concrete examples). No boundary constraints are applied to the reference cells, which have the same initial conditions as the physical cells. The top and bottom edges of the tissue are free boundaries: no external force or displacement is applied to these cells, so they evolve solely due to Equations (6) and (7).

### 2.4. Model parameters

For reference, Table 2 summarises all model parameters, their physical interpretations, their dimensional and dimensionless forms (where applicable), and the values or ranges used in the parameter sweeps of Section 3. Note we restrict our focus here to *α ∈* [0, 0.5] as this represents where *k_R_ ≤ k_S_* and we expect the remodelling force to be weaker than the restoring force but this could be relaxed in future work. We consider *η ∈* [0.01, 1], as this range is sufficient to capture the qualitative behaviours of the model.

**Table 2:**
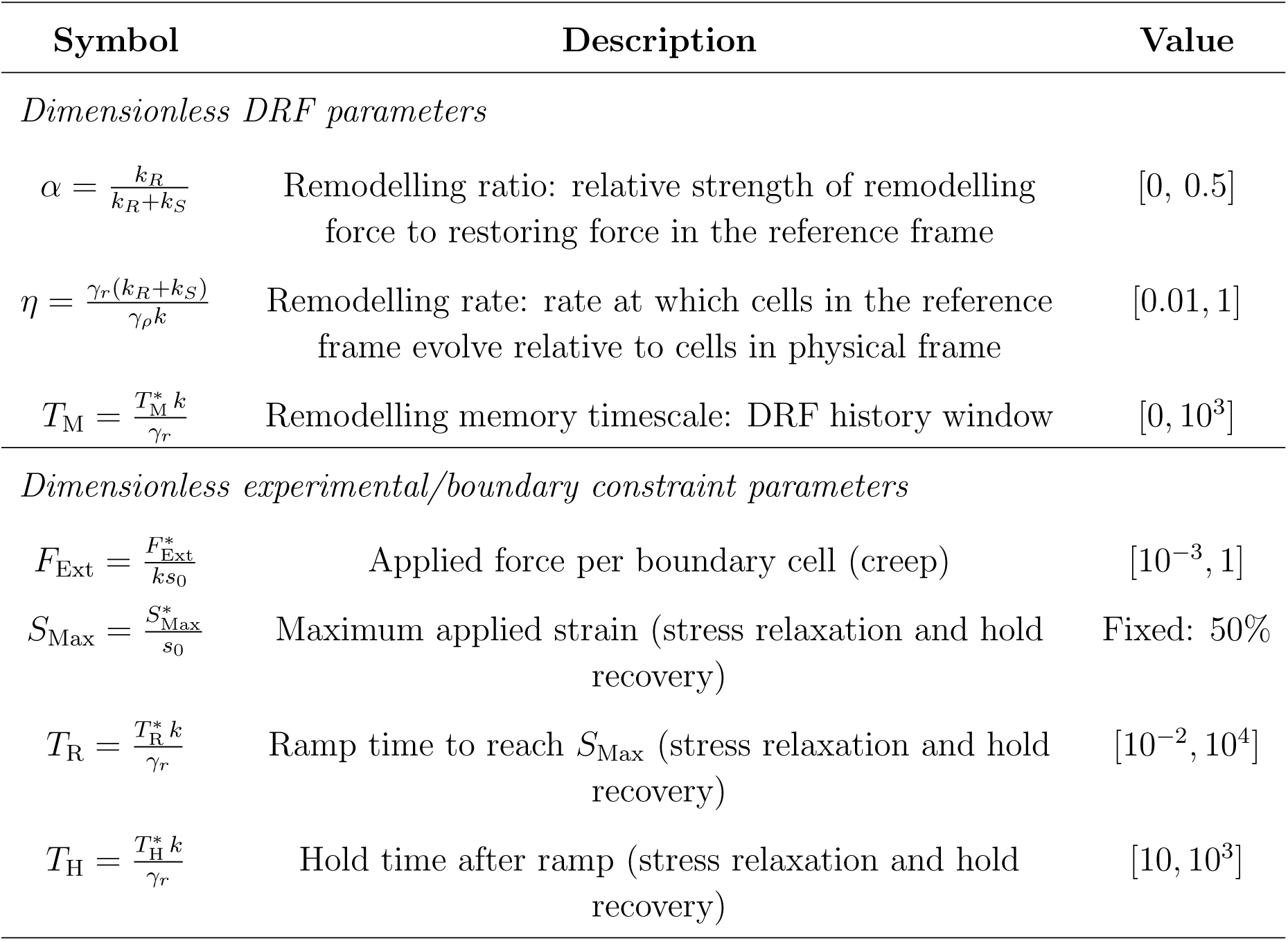
Model parameters. Dimensionless parameters appear in Equations (6), (7) and (8) and are obtained via the non–dimensionalisation, Equation (5). The length scale is *s*_0_ (static cell–cell separation), and the intrinsic timescale is *γ_r_/k* (physical–cell relaxation time). Where a parameter is fixed throughout all simulations, its fixed value is given; otherwise the range explored in the parameter sweeps is stated. Unless specifically stated otherwise, an *∗* denotes the dimensional form of a parameter where a dimensional form is not specifically specified.

| Symbol | Description | Value |
| --- | --- | --- |
| *Dimensionless DRF parameters* |  |  |
| $\alpha = \frac{k_R}{k_R + k_S}$ | Remodelling ratio: relative strength of remodelling force to restoring force in the reference frame | $[0, 0.5]$ |
| $\eta = \frac{\gamma_r(k_R + k_S)}{\gamma_\rho k}$ | Remodelling rate: rate at which cells in the reference frame evolve relative to cells in physical frame | $[0.01, 1]$ |
| $T_M = \frac{T_M^* k}{\gamma_r}$ | Remodelling memory timescale: DRF history window | $[0, 10^3]$ |
| *Dimensionless experimental/boundary constraint parameters* |  |  |
| $F_{\text{Ext}} = \frac{F_{\text{Ext}}^*}{k s_0}$ | Applied force per boundary cell (creep) | $[10^{-3}, 1]$ |
| $S_{\text{Max}} = \frac{S_{\text{Max}}^*}{s_0}$ | Maximum applied strain (stress relaxation and hold recovery) | Fixed: 50% |
| $T_R = \frac{T_R^* k}{\gamma_r}$ | Ramp time to reach $S_{\text{Max}}$ (stress relaxation and hold recovery) | $[10^{-2}, 10^4]$ |
| $T_H = \frac{T_H^* k}{\gamma_r}$ | Hold time after ramp (stress relaxation and hold recovery) | $[10, 10^3]$ |

### 2.5. Implementation details

Once a tissue (and its initial configuration) has been defined and boundary constraints specified, the dimensionless model equations, Equations (6) and (7), are solved numerically using the Matlab ODE solver ode15s, a variable–order solver based on numerical differentiation formulae (NDFs). We use this solver because the model contains two coupled subsystems — for the physical and reference cells — that may evolve on potentially well–separated timescales, rendering the system stiff when *η ≪* 1 (or *η ≫* 1). Note that for *T*_M_ *>* 0, the integral in Equation (8) is approximated using the trapezoidal rule, using the solution at previous time steps. Code for solving the governing equations (and fitting series of exponential functions to the resulting time–series as detailed in Sections 3.1.2 and 3.2.2) can be found, released under an open–source licence, at https://github.com/jmosborne/CellRemodelling.

## 3. Results

We use the DRF model to perform *in silico* versions of creep and stress relaxation experiments, and analyse the timescales of the resulting responses.

### 3.1. Subcellular remodelling alters the relaxation dynamics in creep relaxation experiments

We first consider creep relaxation experiments, where a constant force is applied to the right–hand boundary of the tissue while the left–hand boundary is held fixed.

#### 3.1.1. Creep relaxation experiments

To perform a creep relaxation experiment, we initialise the tissue as described in Section 2.3 and apply a constant force of magnitude *F*_Ext_ (in the positive *x*–direction) to each cell on the right–hand boundary, while holding cells on the left–hand boundary fixed. Specifically, we set 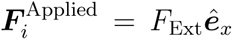 for all cells on the right–boundary and 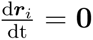 for all cells on the left–boundary (See Figure 2 (c)). We record the strain (i.e., current tissue extension divided by the initial width, indicated in Figure 2 (c)) of the monolayer for both the physical and reference states during the resulting deformation (Figure 2 (d)). Results from two typical creep relaxation simulations are presented in Figure 3, one without remodelling (*F*_Ext_ = 1*, α* = 0) and one with remodelling (*F*_Ext_ = 1*, α* = 0.5*, η* = 0.01).

**Figure 3:**
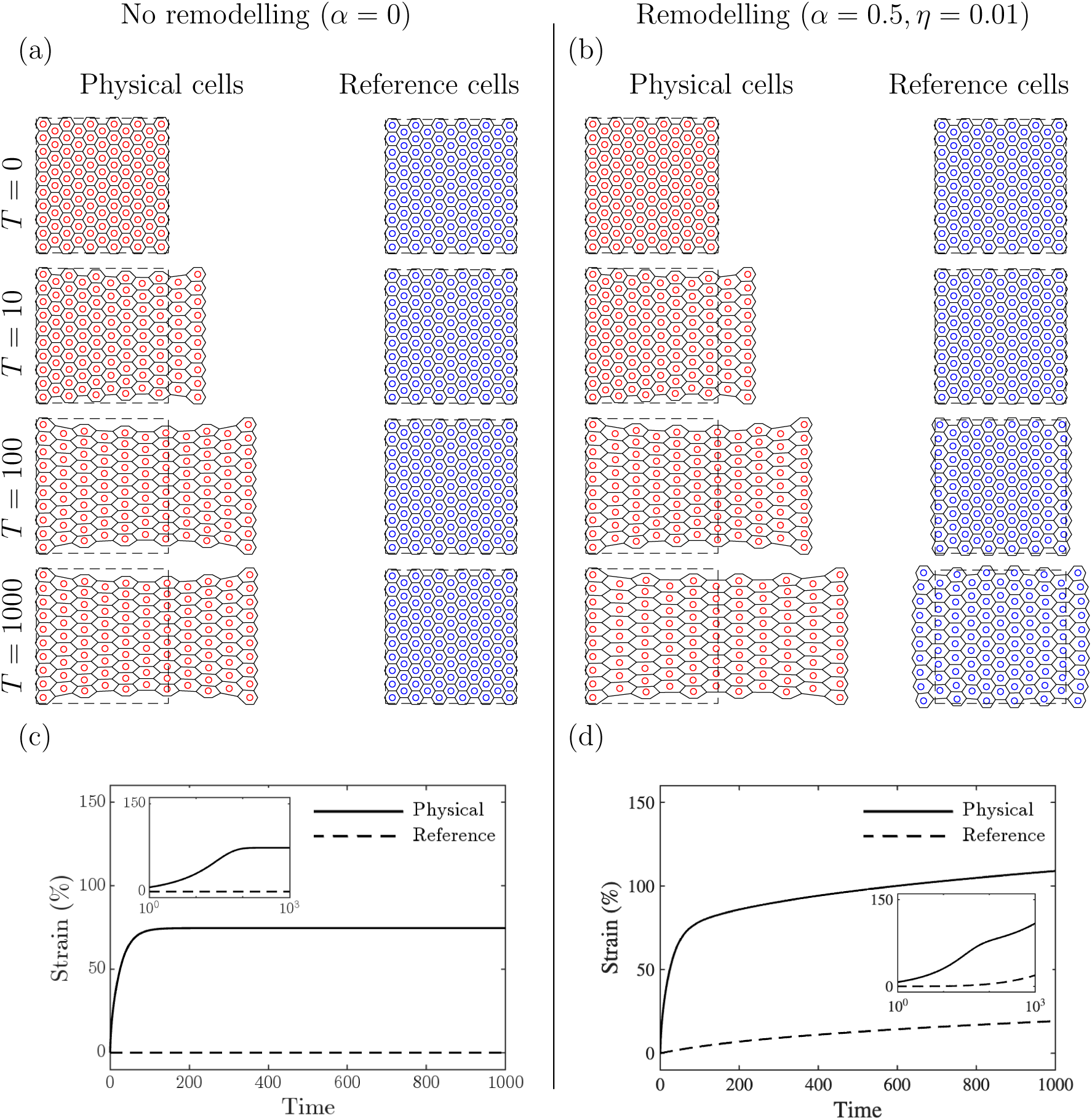
Example creep relaxation experiments. Evolution of the physical and reference state configurations, as an external force, *F*_Ext_ = 1, is applied to cells on the right boundary while cells on the left boundary are held fixed, for (a) no remodelling (*α* = 0) and (b) with remodelling (*α* = 0.5*, η* = 0.01). The dashed outline denotes the initial configuration of the monolayer. Strain of the physical (solid line) and reference (dashed line) states over time for (c) no remodelling and (d) with remodelling. The insets in (c) and (d) show zooms of the main panels plotted on a log time scale to highlight the short–time behaviour. Note *T*_M_ = 0.

Figures 3 (a) and (b) provide snapshots of simulations without and with remodelling; results at later times show how remodelling allows greater deformation of the physical cells. Figures 3 (c) and (d) show the key qualitative difference: without remodelling the tissue experiences a single deformation followed by a plateau. With remodelling, the initial response is the same; however, the strain undergoes an additional, slower deformation. Figure 3 (b) shows that the slower strain dynamics are driven by continued evolution of the reference frame when remodelling is included.

#### 3.1.2. Quantifying relaxation timescales

We have shown that including a dynamic reference frame changes the cells’ response to creep relaxation experiments. We now characterise these dynamics in a phenomenological manner by fitting sums of up to 10 exponentials to time–series data extracted from our simulation results. The functional form used to fit these simulation data is motivated by the observation that the solution to a highly simplified version of this model (i.e., a one–dimensional chain of linear springs) is a sum of exponentials [35].

Figure 4 illustrates the procedure used to fit simulated strain response curves. Let *ε*(*t*) denote the simulated strain at time *t*, defined at a number of computational time–steps (shown in Figure 4 (a) and (b)). To compare simulations across different applied forces and parameter values, we proceed as follows.

i. Truncate each time–series at the first equilibrium time *t* = *T*_End_, defined by the condition 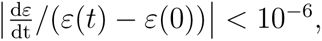,§ (see the red line in Figure 4 (a) and (b)) then rescale both strain and time to [0, 1].

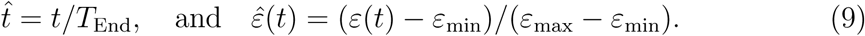
ii. The rescaled strain *ε*^(*t*) is sampled at *N* = 1001, || log–uniformly spaced time points *t_i_ ∈* (0, 1] to give values *y_i_* = *ε*^(*t_i_*) (see the black dots in Figure 4 (c) and (d)).
iii. The re–sampled points are then fitted with a truncated exponential series

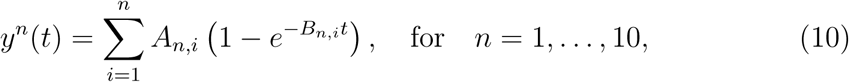

where *A_n,i_*and *B_n,i_*, for *i* = 1*, … , n*, are the fitting parameters (coefficient and decay rate respectively), associated with the functions *y^n^*(*t*) for *n* = 1*, … ,* 10. The fits are calculated using non–linear least squares [36]. The resulting fits are presented in Figure 4 (c) and (d) and the corresponding coefficients in (e) and (f) (see also Table 3).
iv. We define the root mean square error of each fit to be

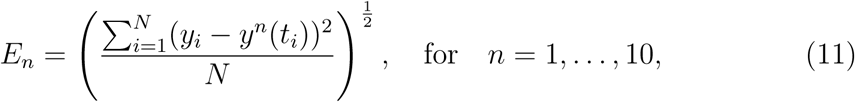

where *y_i_*is the value of the response curve at the *i*th re–sampled time point, *t_i_*. This error, along with the Akaike Information Criterion (AIC) [37] for each fit, is shown in Figure 4 (g) and (h) (see also Table 3).
v. We use an error threshold *E*_T_ = 0.05,¶ to classify the strain response: if *E*_1_ *< E*_T_ the single–exponential fit is the best parsimonious description and the response is dominated by a single timescale; if *E*_1_ *≥ E*_T_, multiple timescales are required to capture the behaviour. (See the dashed lines in Figure 4 (g) and (h), insets).

**Figure 4:**
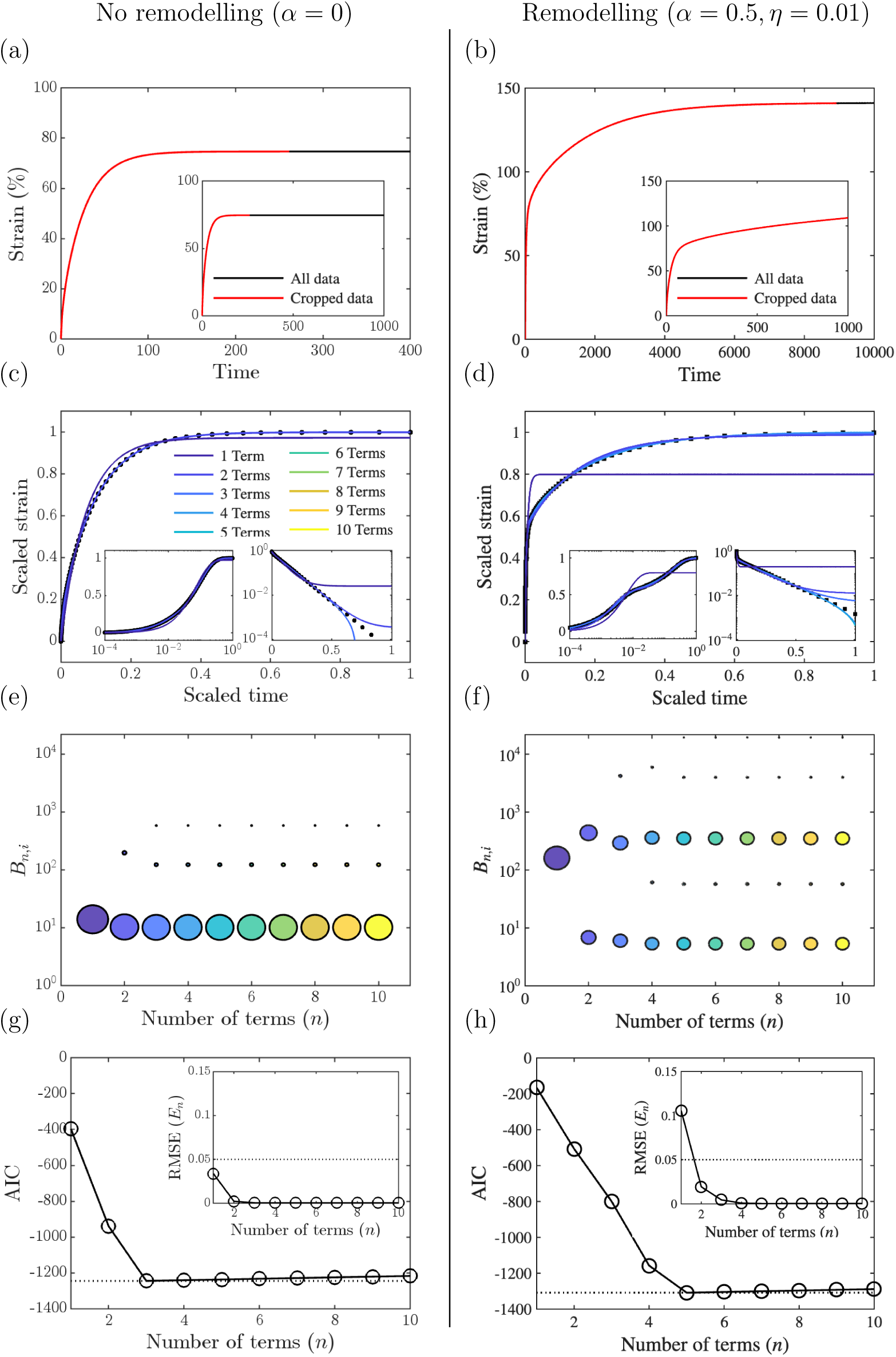
Examples and properties of exponential fits for creep experiments. (left) Creep experiment with no remodelling (*α* = 0.0, *F*_Ext_ = 1, see Figure 3 (a) and (c)). (right) Creep experiment with remodelling (*α* = 0.5, *η* = 0.01, *F*_Ext_ = 1, see Figure 3 (b) and (d)). (a) and (b) show the strain associated with each simulation. The equilibrium segment of the time–series, where the scaled rate of change of the strain is less than 10*^−^*^6^ is indicated in black and the non–equilibrium part (used for rescaling) is indicated in red. The inset shows the time–series on the same scale as Figure 3. (c) and (d) show the re–scaled strain time–series and the corresponding fits with *n* = 1*, … ,* 10 exponentials. Insets show the strain and exponential fits on a log timescale (left) and the 1*−*strain time–series and exponential fits with a log scale on the *y*–axis to highlight the error of the fits (right). (e) and (f) show the coefficients of the exponential fits. The centre of each circle represents the exponential decay rate, *B_n,i_* (for *i* = 1*, … , n*), and the radius represents the coefficient, *A_n,i_*(for *i* = 1*, … , n*), using the same color scheme as (c) and (d). (g) and (h) show the Akaike Information Criterion (AIC) for each fit, with a dashed line at the minimum value; insets show the root mean square error of exponential fits, *E_n_*, for *n* = 1*, … ,* 10, with a dashed line at the threshold value *E_n_*= 0.05. Note *T*_M_ = 0.

**Table 3:** Least squares error (*E_n_*) and the constant parameters in the exponential fits (*A_n,i_, B_n,i_*, with the decay rates *B_n,i_*in brackets) for the creep relaxation experiment, for simulations (a) with no remodelling and (b) with remodelling (see Equations (10) and (11)). Data are taken from the simulations and exponential fits presented in Figure 4. Errors in bold are those above the threshold of *E*_T_ = 0.05.

| (a) Creep relaxation without remodelling ($\alpha = 0.0$, $F_{\text{Ext}} = 1$) | | | | |
| --- | --- | --- | --- | --- |
| $n$ | 1 | 2 | 3 | 4 |
| $E_n$ | $3.3716 \times 10^{-2}$ | $2.2687 \times 10^{-3}$ | $4.9440 \times 10^{-4}$ | $4.9440 \times 10^{-4}$ |
| $A_{n,1}$ | $9.7340 \times 10^{-1}$ | $8.7763 \times 10^{-1}$ | $8.6690 \times 10^{-1}$ | $8.6690 \times 10^{-1}$ |
| $(B_{n,1})$ | $(1.3895 \times 10^1)$ | $(1.0250 \times 10^1)$ | $(1.0066 \times 10^1)$ | $(1.0066 \times 10^1)$ |
| $A_{n,2}$ | | $1.2204 \times 10^{-1}$ | $1.0372 \times 10^{-1}$ | $8.9450 \times 10^{-2}$ |
| $(B_{n,2})$ | | $(1.9667 \times 10^2)$ | $(1.2241 \times 10^2)$ | $(1.2241 \times 10^1)$ |
| $A_{n,3}$ | | | $3.0110 \times 10^{-2}$ | $1.2740 \times 10^{-2}$ |
| $(B_{n,3})$ | | | $(5.8219 \times 10^2)$ | $(1.2242 \times 10^2)$ |
| $A_{n,4}$ | | | | $3.0111 \times 10^{-2}$ |
| $(B_{n,4})$ | | | | $(5.8218 \times 10^2)$ |

Table 3: Least squares error ( $E_n$ ) and the constant parameters in the exponential fits ( $A_{n,i}$ , $B_{n,i}$ , with the decay rates $B_{n,i}$ in brackets) for the creep relaxation experiment, for simulations (a) with no remodelling and (b) with remodelling (see Equations (10) and (11)).
| (b) Creep relaxation with remodelling ($\alpha = 0.5$, $\eta = 0.01$, $F_{\text{Ext}} = 1$) | | | | |
| --- | --- | --- | --- | --- |
| $n$ | 1 | 2 | 3 | 4 |
| $E_n$ | **$1.0574 \times 10^{-1}$** | $1.9319 \times 10^{-2}$ | $4.4814 \times 10^{-3}$ | $7.5568 \times 10^{-4}$ |
| $A_{n,1}$ | $7.9907 \times 10^{-1}$ | $4.5887 \times 10^{-1}$ | $4.4058 \times 10^{-1}$ | $4.1779 \times 10^{-1}$ |
| $(B_{n,1})$ | $(1.6131 \times 10^2)$ | $(6.8540 \times 10^0)$ | $(6.0188 \times 10^0)$ | $(5.3481 \times 10^0)$ |
| $A_{n,2}$ | | $5.2867 \times 10^{-1}$ | $4.6906 \times 10^{-1}$ | $8.6094 \times 10^{-2}$ |
| $(B_{n,2})$ | | $(4.3941 \times 10^2)$ | $(2.9352 \times 10^2)$ | $(6.1184 \times 10^1)$ |
| $A_{n,3}$ | | | $8.5585 \times 10^{-2}$ | $4.3135 \times 10^{-1}$ |
| $(B_{n,3})$ | | | $(4.2235 \times 10^3)$ | $(3.6133 \times 10^2)$ |
| $A_{n,4}$ | | | | $6.6240 \times 10^{-2}$ |
| $(B_{n,4})$ | | | | $(5.9490 \times 10^3)$ |

Inspection of the exponential fits (Figures 4 (c)–(f)) and the corresponding AIC values and errors (Figures 4 (g) and (h)), together with Table 3, reveals that without remodelling *E*_1_ = 0.0337 *< E*_T_, so the response is best described by a single timescale. The AIC selects *n* = 3 as optimal, but the two additional terms have small coefficients (Table 3), arising from the spatial propagation of forces across multiple cells rather than from a distinct physical process; at most three non–zero terms are identified regardless of how many are included in the fit (Figure 4 (e)). With remodelling, the single exponential is no longer the most parsimonious description: *E*_1_ = 0.1057 *> E*_T_, confirming that the response requires multiple timescales. The AIC selects *n* = 5 as the best fit (Figure 4 (f)), and at most five non–zero terms are identified. The two dominant timescales correspond to the fast elastic response of the physical cells and the slower evolution of the reference frame respectively (compare Figures 3 (c) and (d)).

#### 3.1.3. Interpreting creep relaxation experiments

We now investigate how the DRF parameters influence the strain response. Figure 5 shows how the dynamics change as we vary *α ∈* [0, 0.5] and *η ∈* [0.01, 1]. Time–series of the responses and corresponding exponential fits, for varying *α* and *η*, are given in Figures 5 (a) and (d) respectively. The coefficients and errors of the exponential fits are given in Figures 5 (b)–(c) and (e)–(f) respectively. The errors associated with the single exponential fits are presented in Figure 5 (g); in blue regions the dynamics can be approximated by a single timescale (*E*_1_ *<* 0.05) whereas in red regions, they evolve on multiple timescales (*E*_1_ *>* 0.05).

**Figure 5:**
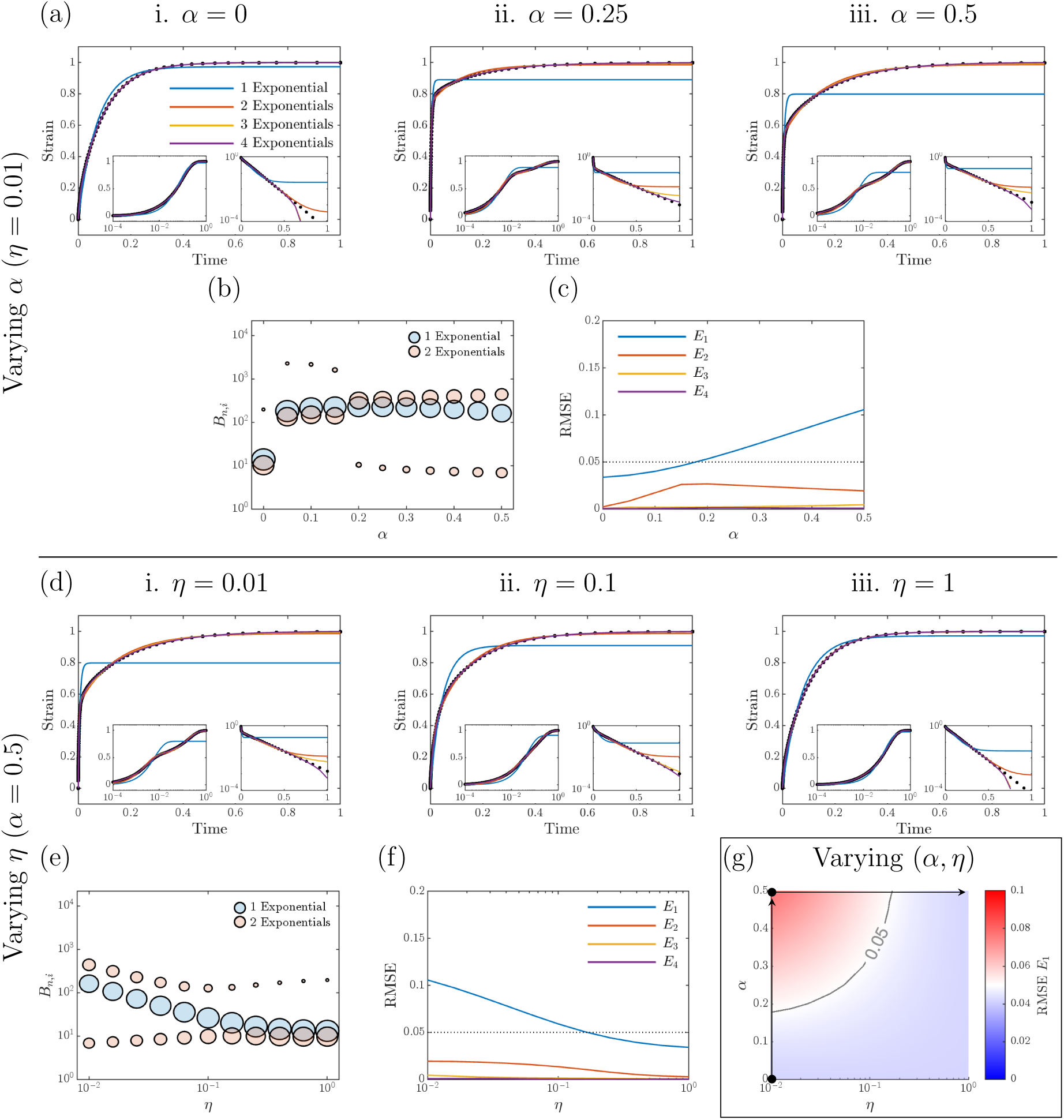
Variation of the dynamic parameters (*α* and *η*) leads to the emergence of multiple timescales in creep relaxation experiments. (a) and (d) show how the re–scaled strain of the tissue evolves under an applied force *F*_Ext_ = 1, as *α* and *η* are varied, respectively. (b) and (e) show the coefficients of the one (blue) and two (red) exponential fits as *α* and *η* are varied, respectively. (c) and (f) show the root mean square error of the exponential fits as *α* and *η* are varied, respectively, when *F*_Ext_ = 1. (g) shows the root mean square error of a single exponential fit for *F*_Ext_ = 1 as *η* and *α* are varied. The arrows in (g) represent the single–variable sweeps presented in (a)–(c) and (d)–(f). Dotted black lines in (c) and (f) and the solid black line in (g) show *E*_1_ = 0.05, which we use as a threshold to indicate an acceptable fit. Note *T*_M_ = 0.

Figure 5 shows that the creep response is well approximated by a single timescale when *α* is small or *η* is large, and that multiple timescales emerge when *α* is increased or *η* is decreased. Mechanistically, increasing *α* makes the DRF more sensitive to the physical state, while decreasing *η* increases the timescale on which the DRF evolves relative to the timescale for evolution of the physical cells. The interface (in parameter space) between single– and multiple–timescale behaviour (*E*_1_ = 0.05, shown in Figure 5 (g)) does not depend on the magnitude of the applied force (results not shown for brevity; however, this is predicted by the constant errors for varying *F*_Ext_ shown in Figure 6 (a) below).

**Figure 6:**
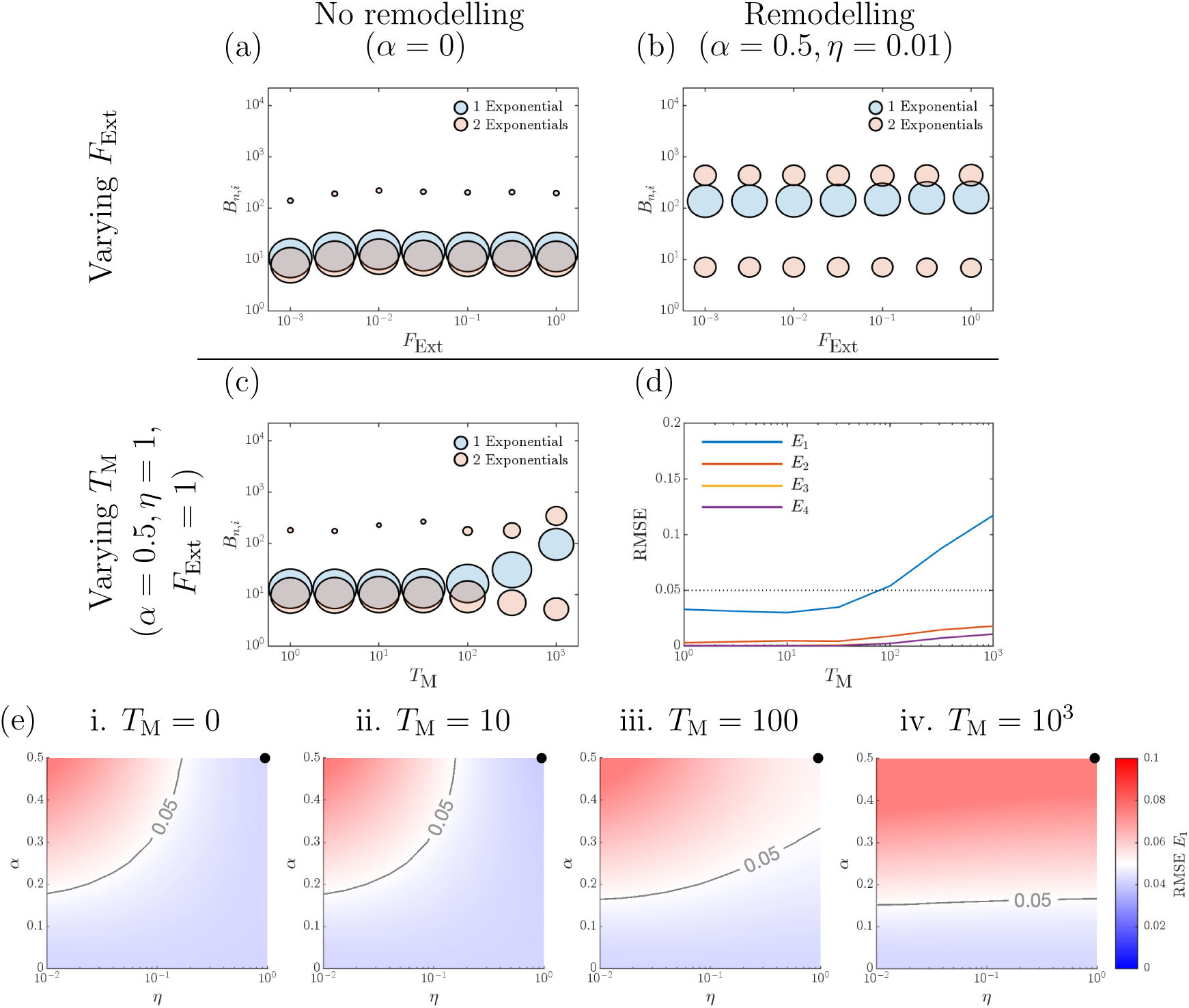
The number of timescales in creep experiment is independent of the applied force. (a) and (b) show how the coefficients of the one (blue) and two (red) exponential fits vary as the applied force, *F*_Ext_, is varied, without (*α* = 0.0) and with remodelling (*α* = 0.5*, η* = 0.01) respectively. All simulations in (a) and (b) are with no memory, *T*_M_ = 0. Example time–series, exponential fits, and error plots are given in Supplementary Figure S1. **Memory leads to multiple timescales in creep experiment.** (c) and (d) show the coefficients of the one (blue) and two (red) exponential fits, and root mean square error of the exponential fits as *T*_M_ is varied for *α* = 0.5 and *η* = 1. (e) shows the root mean square error of single exponential fit as *η* and *α* are varied for increasing memory *T*_M_. The black circles denote the values of (*α, η*) used in (c) and (d). Example time–series and exponential fits are given in Supplementary Figure S2. Dotted black line in (d) and solid black line in (e) show the threshold error of *E*_T_ = 0.05.

In Figures 6 (a) and (b), we show how the coefficients of the exponential fits vary as the applied force, *F*_Ext_, is increased for cases with no remodelling (*α* = 0) and with remodelling (*α* = 0.5*, η* = 0.01). Time–series of the responses, exponential fits, and error plots are given in Supplementary Figure S1. In both cases, and for all parameter sets (*α, η*), the magnitude of the applied force does not affect the shape of the strain response. Further, the coefficients (and therefore errors) are approximately constant as *F*_Ext_ varies. This suggests that the DRF parameters, rather than the applied force, influence the behaviour of the strain response.

Cell remodelling due to imposed stress or strain may depend on the loading history, rather than on instantaneous values. In our model, this is captured by the history–dependent term in Equation (8). We now investigate how the dynamics of the strain response change as we vary the remodelling memory (*T*_M_ *>* 0). Figures 6 (c) and (d) show how the coefficients and errors of the exponential fits of the strain response change as *T*_M_ varies when *α* = 0.5 and *η* = 1 (shown by black dots in Figure 6 (e)), which experiences a single timescale response when *T*_M_ = 0. Time–series of the response and exponential fits are given in Supplementary Figure S2. As *T*_M_ is increased, the error for the one exponential fit increases (Figure 6 (d)) and the coefficients of the 2–exponential fits transition from being dominated by one term (for small *T*_M_) to both terms being significant (large *T*_M_), Figure 6 (c).

Figure 6 (e) shows how the remodelling memory, *T*_M_, affects the transition between single and multiple timescale responses (here defined, without loss of generality, to be the *E*_1_ = 0.05 nullcline). As *T*_M_ increases the number of parameter sets (*α, η*) characterised by a multiple timescale strain response also increases. The remodelling memory can therefore introduce additional timescales into a system which might otherwise be characterised by a single timescale response. We note that this result is independent of the applied forces as already shown for *T*_M_ = 0 (results not shown for brevity).

### 3.2. Subcellular remodelling alters the relaxation dynamics in stress relaxation experiments

We now investigate how the DRF changes the system’s response to stress relaxation experiments.

#### 3.2.1. Stress relaxation experiments

We simulate stress relaxation experiments by holding cells on the left–hand edge of the tissue fixed 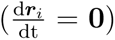 and prescribing the displacement of cells on the right–hand edge (***r****_i_*(*t*) = ***r***_Ext_(*t*)). A constant strain rate is applied until the strain in the monolayer increases to a prescribed maximum, *S*_Max_ (here we set *S*_Max_ = 50%, for consistency with the range of strain values measured by Harris et al. [6]). The time *T*_R_ for the monolayer to attain the maximum strain is termed the *ramp–time*. The monolayer is then held at the constant strain of *S*_Max_ = 50% for the *hold–time*, *T*_H_. We record the stress at the right–hand edge of the monolayer during the resulting deformation (i.e., for *T*_R_ *< t ≤ T*_R_ + *T*_H_). This is denoted by *σ*(*t*) and is calculated by averaging the resistive forces on the right–hand edge of the monolayer, *F*_Res_ (see Figure 2 (c)), and dividing by the length of the edge.

Figures 7 (a) and (b) provide snapshots of simulations without and with remodelling. Figures 7 (c) and (d) show how the stress and strain evolve during the ramp–time (0 *≤ t ≤ T*_R_) and the hold–time (*T*_R_ *< t ≤ T*_R_ + *T*_H_). In both cases, the stress, *σ*(*t*), converges to an equilibrium stress, *σ_∞_*, which is different in each case. We define *σ_∞_* to be the value of *σ*(*t*) when 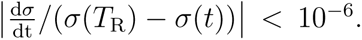.^+^ Figures 7 (e) and (f) show how *σ − σ_∞_* evolves during the hold–time (the insets provide detailed information on a log scale).

**Figure 7:**
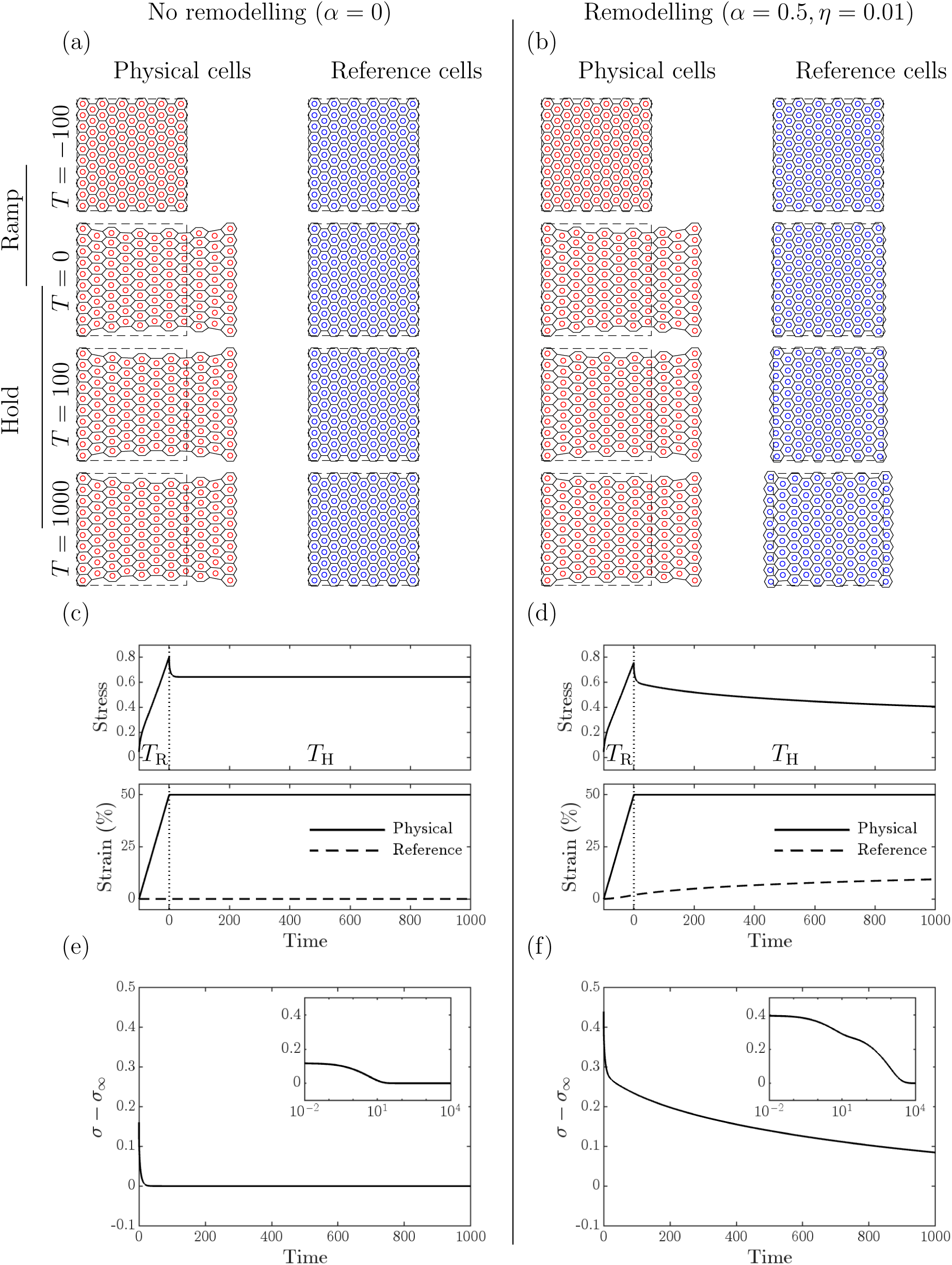
Example stress relaxation simulations. Physical and reference state configurations over time, as an external strain is applied (then subsequently held, *T*_R_ = 100 and *T*_H_ = 1000) to cells on the right edge while cells on the left edge are kept fixed, for (a) no remodelling (*α* = 0) and (b) with remodelling (*α* = 0.5*, η* = 0.01). The dashed outline denotes the initial monolayer. Stress and strain distribution over time for (c) no remodelling and (d) with remodelling. Stress distribution after the strain ramp for (e) no remodelling and (f) with remodelling. Insets in (e) and (f) show the stress using a log scale on the *x*–axis to highlight the short–time behaviour. Note *T*_M_ = 0.

As for the creep experiment, when remodelling is neglected, relaxation occurs on a single timescale during the hold phase, while the tissue evolves to an equilibrium plateau (Figure 7 (e)). By contrast, when remodelling is included, we observe an additional relaxation period (see Figure 7 (f)). This relaxation phase is due to the evolution of the reference frame during the simulation (compare Figures 7 (a) and (b), and the dashed lines in Figures 7 (c) and (d)).

#### 3.2.2. Quantifying relaxation timescales

As for the creep relaxation experiments, we identify different timescales during the stress relaxation experiments on which different dynamics occur. In order to infer the number of timescales in the response, we again fit the stress time–series data with sums of exponentials.

Figure 8 shows the procedure used to compare stress relaxation simulation time–series. We analyse the stress relaxation time–series using the same (i) truncation (this time using *σ_∞_*), rescaling, and (ii) log–resampling procedure as for creep relaxation time–series (See Section 3.1.2 (i)–(v) and Figure 8 (a)–(d)), but with a different fitting form in Step (iii) to reflect the decay of the stress signal during the hold phase.

**Figure 8:**
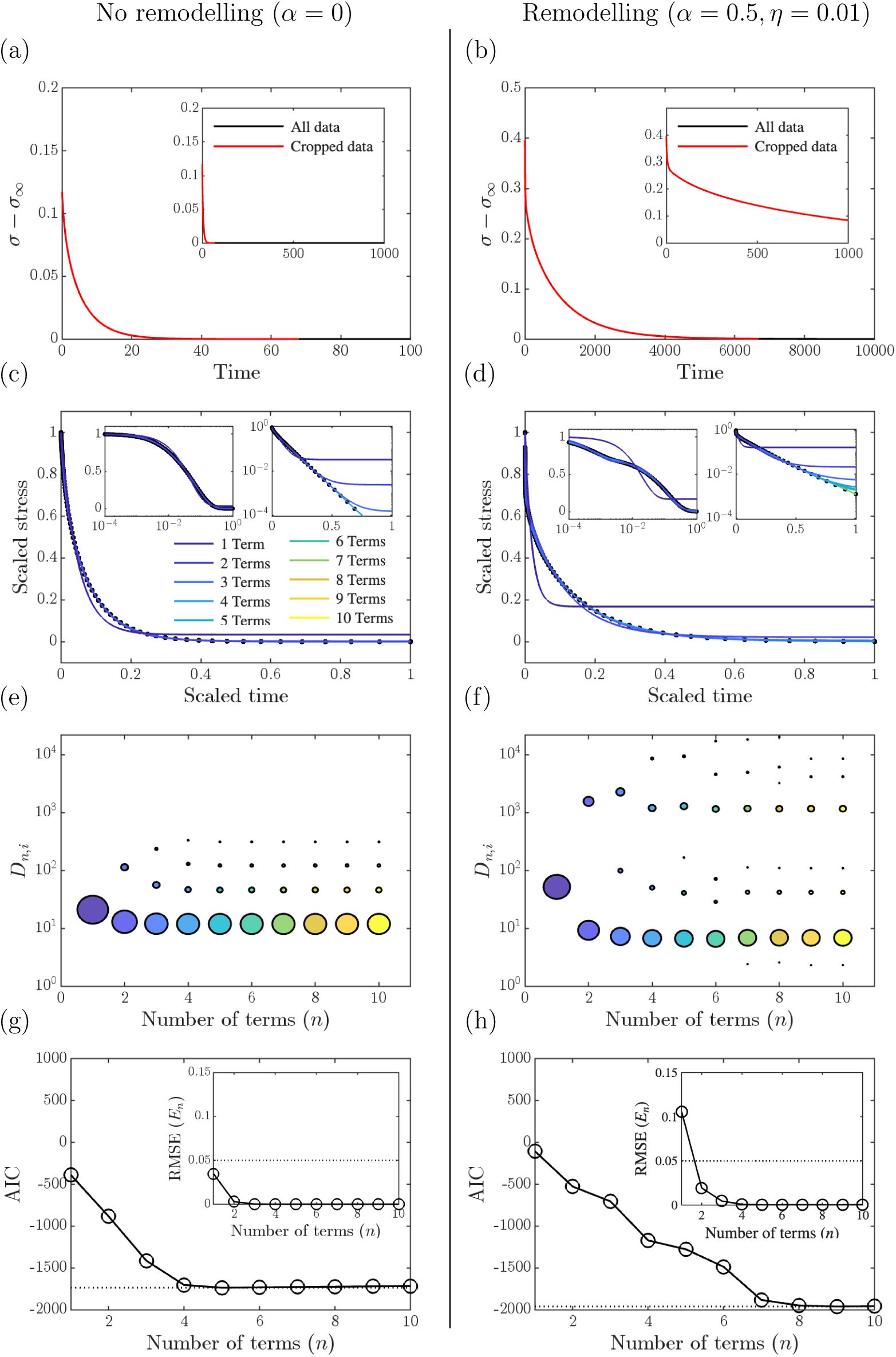
Examples and properties of exponential fits for stress relaxation experiments. (left) Stress relaxation experiment with no remodelling (*α* = 0.0, *T*_R_ = 100, see Figure 7 (a) and (c)). (right) Stress relaxation experiment with remodelling (*α* = 0.5, *η* = 0.01, *T*_R_ = 100, see Figure 7 (b) and (d)). (a) and (b) show the stress of each simulation. The segments of the time–series with a scaled rate of change under 10*^−^*^6^ (which is used to signify equilibrium) are in black and the non–equilibrium part (used for rescaling) is given in red. Insets show the time–series on the same scale as Figure 7. (c) and (d) show the stress time–series mapped and re–scaled to [0, 1] *×* [0, 1] along with fits using *n* = 1*, … ,* 10 exponentials. Insets show the stress time–series and exponential fits on a log timescale (left) and with a log scale on the *y*–axis to highlight the error of the fits (right). (e) and (f) show coefficients of the exponential fits. The centre of each circle represents the exponential decay rate, *D_n,i_*(for *i* = 1*, … , n*), and the radius represents the coefficient, *C_n,i_*(for *i* = 1*, … , n*), using the same color scheme as (c) and (d). (g) and (h) show the Akaike Information Criterion (AIC) for each exponential fit, with a dashed line at the minimum value, insets show the root mean square error of exponential fits, *E_n_*, for *n* = 1*, … ,* 10, with a dashed line at *E_n_*= 0.05. Note *T*_M_ = 0.

(iii) We replace Equation (10) with

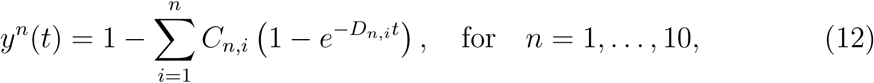

where *C_n,i_*and *D_n,i_*, for *i* = 1*, … , n*, are the fitting parameters (coefficient and decay rate respectively), for the functions *y^n^*(*t*) for *n* = 1*, … ,* 10. The resulting fits are shown as coloured lines in Figures 8 (c) and (d) and the coefficients are shown in Figures 8 (e) and (f) (see also Table 4).
(iv) The root mean square error is calculated as in Equation (11) and is plotted with the AIC for each fit in Figure 8 (g) and (h) (see also Table 4).
(v) We use a threshold of *E*_T_ = 0.05 to determine whether a single exponential is an appropriate parsimonious fit to the data. (See the dashed lines in Figure 8 (g) and (h), insets)

**Table 4:** Fit error (*E_n_*) and exponential fit parameters (*C_n,i_, D_n,i_*, with decay rates *D_n,i_*in brackets) for the stress relaxation experiment; see Equations (11) and (12). Results are shown for (a) no remodelling and (b) remodelling. Data are taken from the simulations and exponential fits presented in Figure 8. Errors in bold are those above the threshold of *E*_T_ = 0.05.

| (a) Stress relaxation without remodelling ($\alpha = 0.0$, $T_R = 100$) | | | | |
| --- | --- | --- | --- | --- |
| $n$ | 1 | 2 | 3 | 4 |
| $E_n$ | $3.4721 \times 10^{-2}$ | $2.9805 \times 10^{-3}$ | $2.1443 \times 10^{-4}$ | $5.0620 \times 10^{-5}$ |
| $C_{n,1}$ | $9.6599 \times 10^{-1}$ | $7.7814 \times 10^{-1}$ | $7.1322 \times 10^{-1}$ | $7.0153 \times 10^{-1}$ |
| $(D_{n,1})$ | $(2.1004 \times 10^1)$ | $(1.2996 \times 10^1)$ | $(1.2022 \times 10^1)$ | $(1.1910 \times 10^1)$ |
| $C_{n,2}$ | | $2.1943 \times 10^{-1}$ | $2.0953 \times 10^{-1}$ | $1.7787 \times 10^{-1}$ |
| $(D_{n,2})$ | | $(1.1383 \times 10^2)$ | $(5.6248 \times 10^1)$ | $(4.6738 \times 10^1)$ |
| $C_{n,3}$ | | | $7.7107 \times 10^{-2}$ | $8.7956 \times 10^{-2}$ |
| $(D_{n,3})$ | | | $(2.3613 \times 10^2)$ | $(1.3005 \times 10^2)$ |
| $C_{n,4}$ | | | | $3.2637 \times 10^{-2}$ |
| $(D_{n,4})$ | | | | $(3.3333 \times 10^2)$ |

Table 4: Fit error ( $E_n$ ) and exponential fit parameters ( $C_{n,i}$ , $D_{n,i}$ , with decay rates $D_{n,i}$ in brackets) for the stress relaxation experiment; see Equations (11) and (12).
| (b) Stress relaxation with remodelling ($\alpha = 0.5$, $\eta = 0.01$, $T_R = 100$) | | | | |
| --- | --- | --- | --- | --- |
| $n$ | 1 | 2 | 3 | 4 |
| $E_n$ | **$1.4254 \times 10^{-1}$** | $1.7486 \times 10^{-2}$ | $7.2010 \times 10^{-3}$ | $7.0590 \times 10^{-4}$ |
| $C_{n,1}$ | $8.3203 \times 10^{-1}$ | $6.5989 \times 10^{-1}$ | $5.9835 \times 10^{-1}$ | $5.6815 \times 10^{-1}$ |
| $(D_{n,1})$ | $(5.2106 \times 10^1)$ | $(9.2615 \times 10^0)$ | $(7.2506 \times 10^0)$ | $(6.7566 \times 10^0)$ |
| $C_{n,2}$ | | $3.1856 \times 10^{-1}$ | $1.3646 \times 10^{-1}$ | $1.3664 \times 10^{-1}$ |
| $(D_{n,2})$ | | $(1.5663 \times 10^3)$ | $(9.9253 \times 10^1)$ | $(5.0430 \times 10^1)$ |
| $C_{n,3}$ | | | $2.5990 \times 10^{-1}$ | $2.1817 \times 10^{-1}$ |
| $(D_{n,3})$ | | | $(2.2823 \times 10^3)$ | $(1.2013 \times 10^3)$ |
| $C_{n,4}$ | | | | $7.5100 \times 10^{-2}$ |
| $(D_{n,4})$ | | | | $(8.6343 \times 10^3)$ |

Inspection of Figures 8 (c) and (d) indicates that a sum of exponentials again provides a good fit to the scaled time–series. The coefficient values for no remodelling in Figure 8 (e) show that, for each fit, there is one dominant exponential term, suggesting a single timescale of interest. When remodelling is active (Figure 8 (d)), there are two dominant terms, suggesting multiple timescales. The AIC (Figures 8 (g) and (h)) shows that the best fit for the no remodelling simulation has *n* = 5 and the best fit for the remodelling simulation has *n* = 9. These values are close to the number of non–zero terms in the fits (Figures 8 (c) and (d)). The errors of the exponential fits reveal that when there is no remodelling the single exponential is a good fit (*E*_1_ = 0.0347) and all other errors are smaller (Figure 8 (g) and Table 4). By contrast, when remodelling is active the single exponential is no longer a good fit (*E*_1_ = 0.1425), and all other errors are smaller, specifically less than 0.05 (Figure 8 (h) and Table 4). Therefore, we again choose to use an error threshold of *E*_T_ = 0.05 to determine whether the time–series can be described by a single relaxation timescale.

#### 3.2.3. Interpreting stress relaxation experiments

We now study the behaviour of the stress response in the stress relaxation experiments.

Figure 9 shows how the stress relaxation dynamics vary with *α ∈* [0, 0.5] and *η ∈* [0.01, 1]. As for the creep experiments, regions with small *α* or large *η* are well approximated by a single timescale, whereas large *α* and small *η* produce multiple–timescale responses. Example time–series and fits are shown in Supplementary Figure S3 (a) and (b).

**Figure 9:**
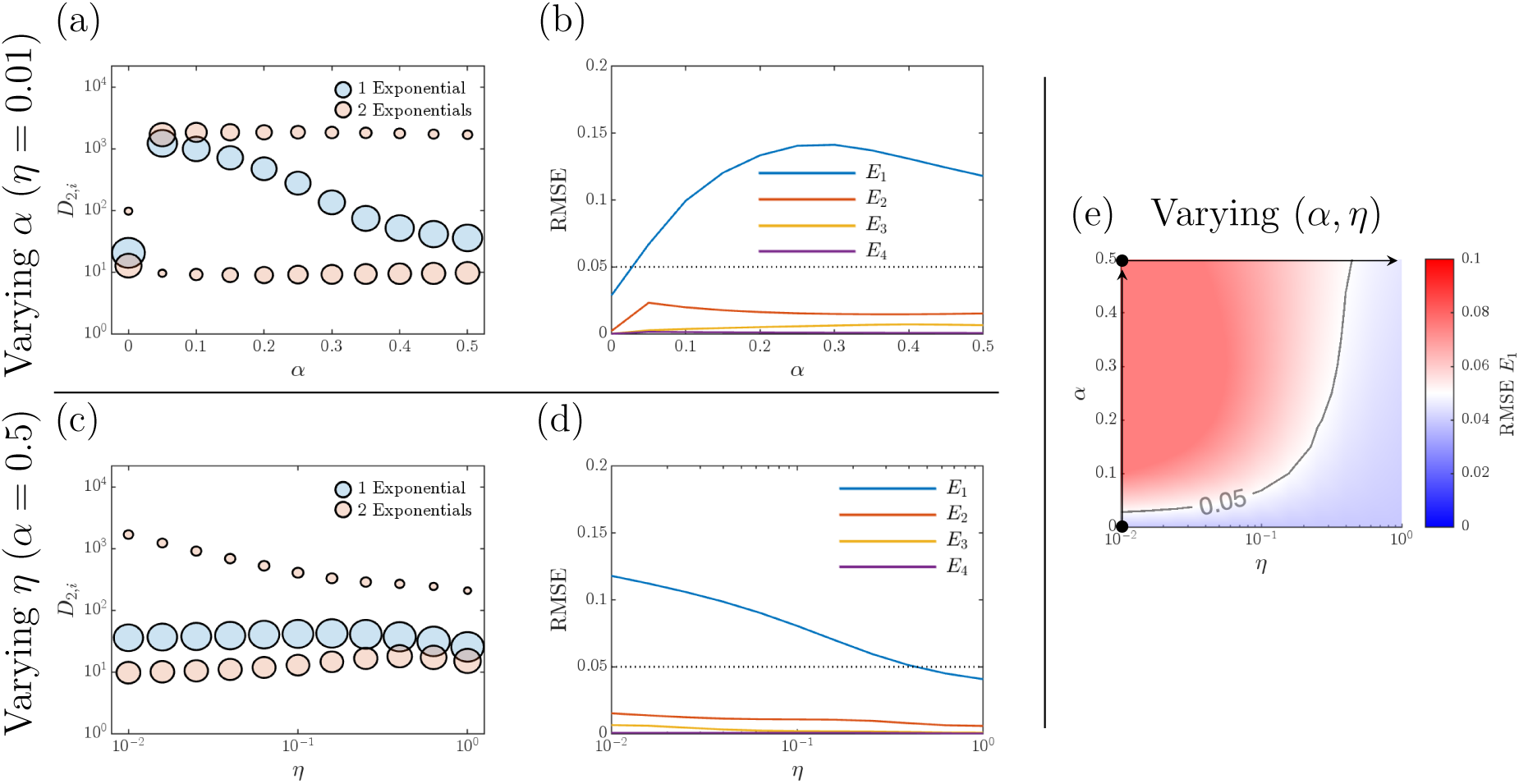
Variation of the dynamic parameters (*α* and *η*) leads to the emergence of multiple timescales in stress relaxation experiments. (a) and (c) show the coefficients of the one (blue) and two (red) exponential fits as *α* and *η* are varied, respectively. (b) and (d) show the root mean square error of the exponential fits as *α* and *η* are varied, respectively. (e) shows the root mean square error of a single exponential fit as *η* and *α* are varied. The arrows in (e) represent the single–variable sweeps presented in (a)–(b) and (c)–(d). All simulations are with *T*_R_ = 100. Example time–series and exponential fits are given in Supplementary Figure S3 (a) and (b). Dotted black lines in (b) and (d) and the solid black line in (e) show an error of *E*_T_ = 0.05, which we use as a threshold to indicate an acceptable fit. Note *T*_M_ = 0.

Figures 10 (a)–(d) show that, unlike in creep experiments, the ramp duration changes the qualitative form of the stress response. Without remodelling the dynamics remain effectively single–timescale, although the fit quality varies with *T*_R_. With remodelling, long ramps allow both the physical and reference states to relax during loading and so favour a single–timescale response, intermediate ramps expose distinct post–ramp relaxation timescales, and very short ramps are dominated by the large initial elastic stress. Thus *T*_R_ acts as an additional control on the number of observable timescales. We now consider non–zero remodelling memory (*T*_M_ *>* 0). Figures 10 (e)–(g) show that increasing *T*_M_ increases the error of the single–exponential fit and expands the region of parameter space exhibiting multiple–timescale stress relaxation.

**Figure 10:**
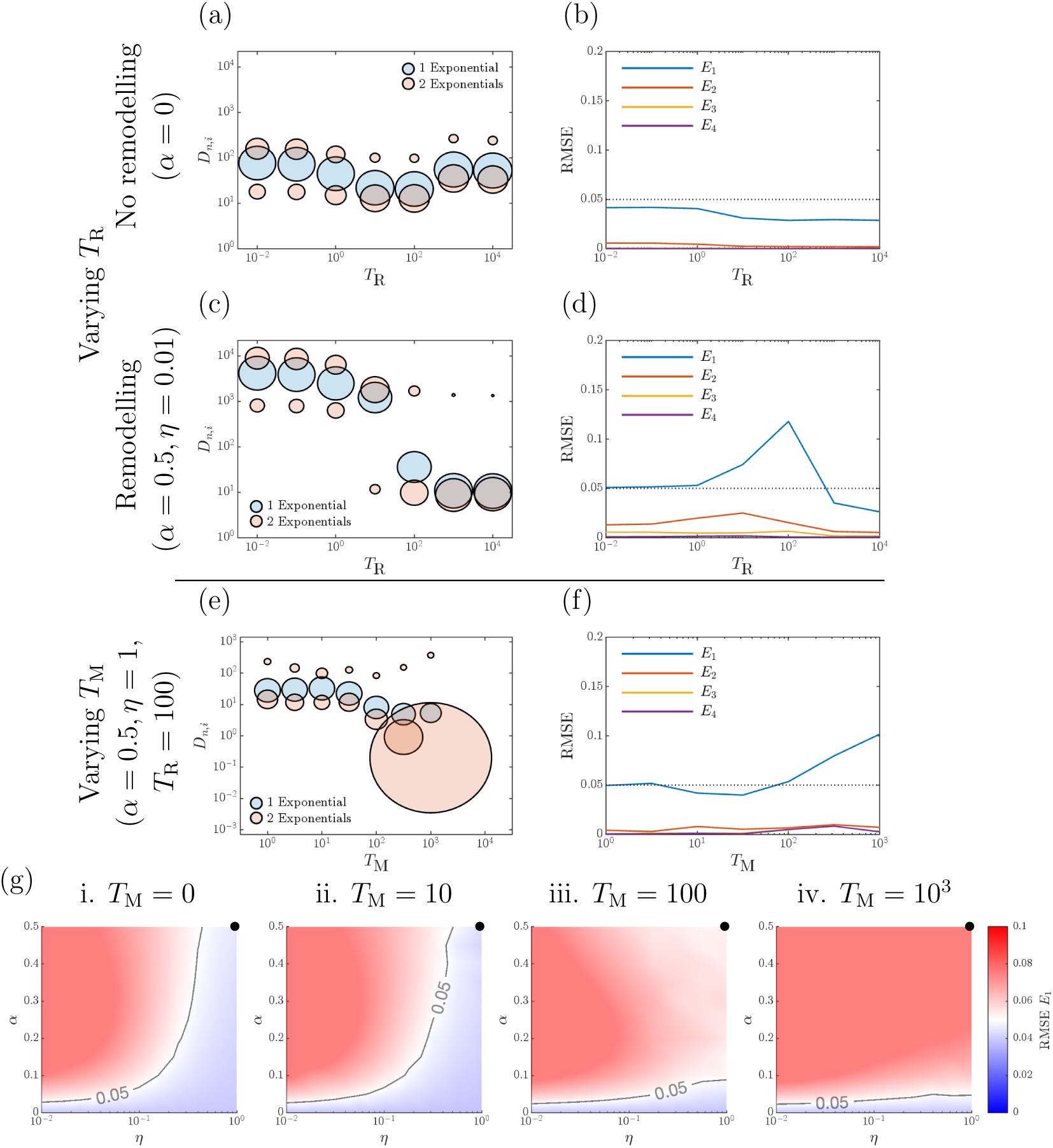
Duration of applied strain can cause multiple timescales in the stress relaxation experiment when remodelling is included. (a) and (c) show how the coefficients of the one (blue) and two (red) exponential fits vary as the ramp–time, *T*_R_, is varied, without (*α* = 0.0) and with remodelling (*α* = 0.5*, η* = 0.01), respectively. (b) and (d) show the root mean square error of the exponential fits as the ramp–time, *T*_R_, is varied without and with remodelling, respectively. Example time–series and exponential fits are given in Supplementary Figure S3 (c) and (d). All simulations in (a)–(d) are with no memory, *T*_M_ = 0. **Memory leads to multiple timescales in the stress relaxation experiment.** (e) and (f) show the coefficients of the one (blue) and two (red) exponential fits, and root mean square error of the exponential fits as *T*_M_ is varied for *α* = 0.5*, η* = 1 and *T*_R_ = 100, which experiences a single timescale response with *T*_M_ = 0. (g) shows the root mean square error of a single–exponential fit as *η* and *α* are varied for *T*_R_ = 100 and increasing memory *T*_M_. The black circles show the values of (*η, α*) used in (e) and (f). Example time–series and exponential fits are given in Supplementary Figure S4. Dotted black lines in (b), (d), and (f) and the solid black line in (g) show the error threshold of *E*_T_ = 0.05.

### 3.3. Monolayer recovery on release from imposed strain depends on duration of imposed strain

A common artefact of many multicellular models is that if a model tissue is held under tension then its relaxation once released is independent of the amount of time for which it was held under tension. This is not the case for real biological tissues, where the rate of strain recovery depends on the time period for which the tissue was deformed [6]. We propose that this discrepancy is due to subcellular remodelling that occurs while the monolayer is held under strain, and the DRF provides a simple framework in which this can be investigated.

We now modify the classical stress relaxation experiment (Section 3.2.1) to describe the above setup and to investigate how the DRF affects tissue dynamics. The tissue is exposed to a strain ramp for *T*_R_ = 100, held for *T*_H_ = 10, 100 or 1000 (i.e., shorter than the ramp time; the same as the ramp time; and longer than the ramp time), and then released. We track the strain relaxation dynamics following release.

In Figure 11, we present snapshots (a)–(c) and strain relaxation responses (d)–(f) for the hold recovery experiment without remodelling, with remodelling, and with both remodelling and non–zero remodelling memory. As expected, in the absence of remodelling the response is independent of *T*_H_ (Figure 11 (d)). By contrast, with remodelling (both with and without memory), the tissue’s rate of strain recovery on release is proportional to 1*/T*_H_ (Figures 11 (e) and (f)). This is because the longer the tissue is held the more the DRF can deform, representing increased subcellular remodelling. To demonstrate this, we consider the stress relaxation experiment from Figure 7 where we hold the tissue at a strain of *S*_Max_ = 50% for *T*_H_ = 10^4^. We then track the stress and strain over time for our three cases. The corresponding stress and strain dynamics (in the cells and in the reference tissue) are presented in Figures 11 (g)–(i). The vertical dashed lines show where a tissue would be released at *T* = 10, 100 and 1000. In Figure 11 (g) the stress in the tissue is the same at *T* = 10, 100 and 1000 so we observe the same response regardless of when the tissue is released (see Figure 11 (d)). By contrast, in Figures 11 (h) and (i), the stress in the tissue decreases from *T* = 10 to *T* = 100 and *T* = 1000 as the strain in the DRF grows, so the longer the tissue is held, the lower the stress at release and the slower the subsequent recovery (see Figures 11 (e) and (f)).

**Figure 11:**
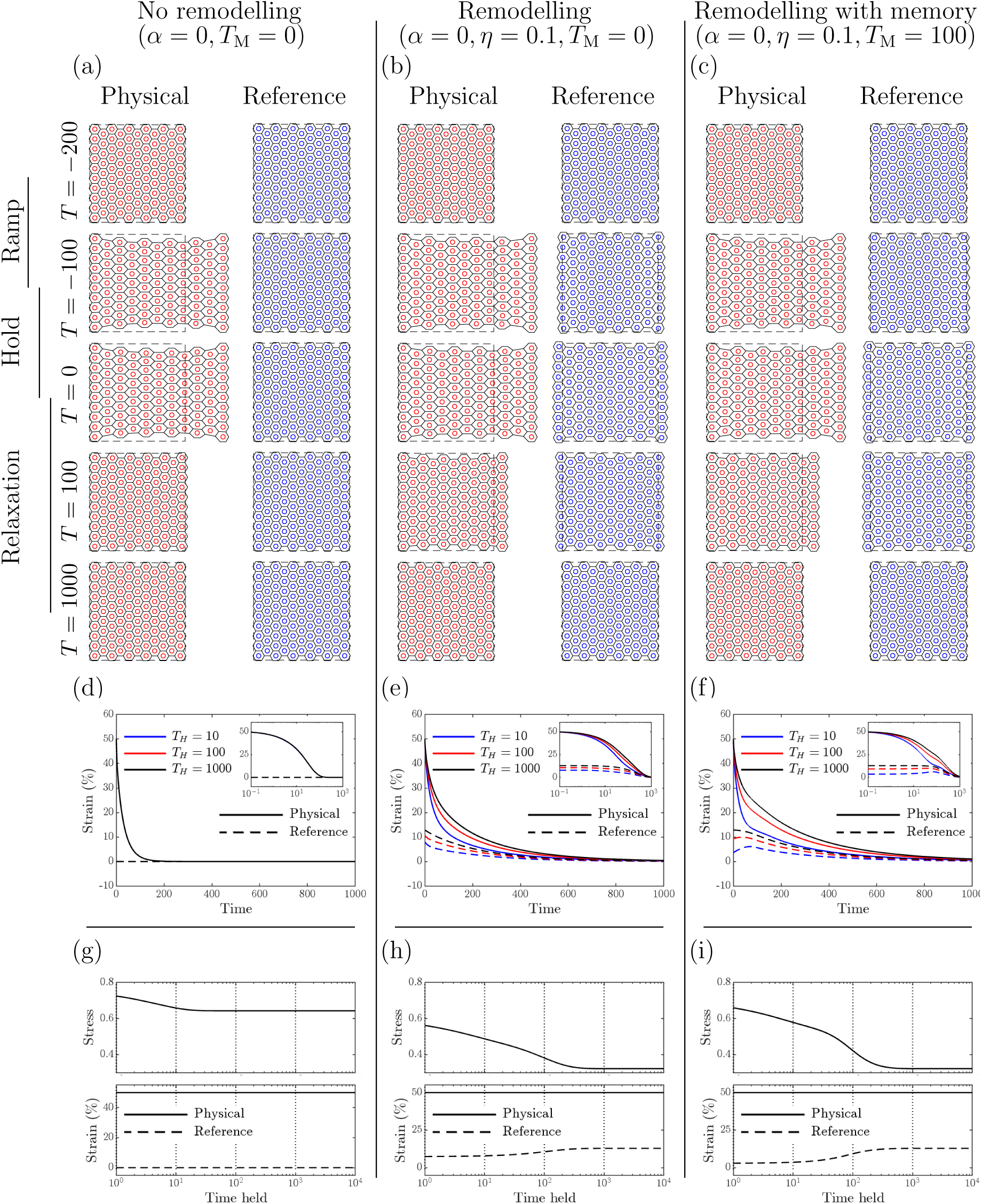
Recovery from deformation depends on duration of deformation. (a)–(c) Physical and reference state configurations over time as an external strain is applied to cells on the right edge (*T*_R_ = 100), subsequently held (*T*_H_ = 100), and then released and allowed to relax, while the left edge is kept fixed, for (a) no remodelling (*α* = 0, *T*_M_ = 0), (b) remodelling (*α* = 0.5, *η* = 0.1, *T*_M_ = 0), and (c) remodelling with memory (*α* = 0.5, *η* = 0.1, *T*_M_ = 100). Strain distributions over time after release for varying hold duration, *T*_H_, are shown for (d) no remodelling, (e) remodelling, and (f) remodelling with memory. Insets show strain distributions on a log timescale. Stress and strain (of the cells and reference frame) over time after an initial application of strain are shown for (g) no remodelling (*α* = 0, *T*_M_ = 0), (h) remodelling (*α* = 0.5, *η* = 0.1, *T*_M_ = 0), and (i) remodelling with memory (*α* = 0.5, *η* = 0.1, *T*_M_ = 100). The vertical dotted lines in (g)–(i) indicate where the tissues in the simulations in (d)–(f) were released from being held.

## 4. Discussion and Conclusion

Experimental results have demonstrated that deformation of epithelial tissues can generate responses which act on multiple timescales; a feature that most existing multicellular models, when formulated with standard elastic interaction laws and without explicit subcellular structure, are unable to capture. We have introduced a dynamic reference frame (DRF) that extends the Voronoi Tessellation model to account for subcellular remodelling, and showed that monolayer recovery on release from imposed strain depends on the duration of that strain. The inclusion of subcellular remodelling (*α >* 0, *η >* 0) introduced additional timescales into the tissue response to creep and stress relaxation experiments, an effect that is more marked when memory is included (*T*_M_ *>* 0). While the model introduces multiple parameters to describe the remodelling, we have shown that the qualitative behaviour of the tissue response is robust to parameter variations, and that the type of response can be controlled by varying the remodelling parameters. These remodelling parameters can be linked to biological processes.

All simulations and parameter sweeps were performed by coupling a DRF to a linear spring, cell centre, Voronoi Tessellation model, arguably one of the simplest multicellular models. However, the framework can be extended to other cell–based models. For example, a DRF could be incorporated into a vertex dynamics model [38], by using a vertex–based reference state.

Our framework can also accommodate changes in the functional forms of the forces linking the physical, reference and static cells, which could be inferred from experimental measurements. Additional intermediate reference states can be introduced to represent further biological processes—such as actin shuttling, cell differentiation, and cell growth—each operating on its own timescale, and the framework extends naturally to three dimensions.

In cell–level models, cell rearrangements due to birth, death, or applied forces require updates to cell–cell connectivity [17]. Since here the reference state shares the same connectivity as the physical state, how it reacts to such changes is paramount if proliferation and death are to be implemented. The simplest approach for including proliferation and death in future work would be to mirror rearrangements in the reference state using techniques from adaptive re–meshing [39], though this would introduce discontinuities into the response and require further work. In addition to changing connectivity, another extension would be to remove the assumption that remodelling parameters are uniform across the tissue, which would allow the study of how heterogeneity in subcellular remodelling affects tissue dynamics.

In conclusion, in this paper, we have presented a new method for studying the effects of subcellular remodelling on deforming tissues, which can be incorporated into multicellular models. The inclusion of a DRF makes it possible to use multicellular models to characterise the dynamic responses of deforming tissues seen experimentally. We anticipate that, through specialisation of the model to specific tissues, this framework will allow more detailed models of organ development and function to be made.

## Conflict of Interest Statement

The authors declare that they have no competing interests.

## Data Access Statement

The code and data required to reproduce the results presented in this study are available, under an open–source licence, at https://github.com/jmosborne/CellRemodelling.

## Ethics Statement

This study involved only mathematical modelling and computational simulations and did not involve human participants, human tissue, or animals. Ethical approval was therefore not required.

## Acknowledgements

The authors gratefully acknowledge the contributions of Alistair Martin and Bayar Menzat to earlier versions of this work, which was developed in part through student projects at the University of Oxford. JMO’s research is supported by the Australian Research Council (ARC) DP23010038, FT230100352, DP260100767. YD’s research is supported by the European Research Council (ERC) under the European Union’s Horizon 2020 research and innovation programme (grant agreement no 803074).

## Supplementary Figures

**Supplementary Figure S1:**
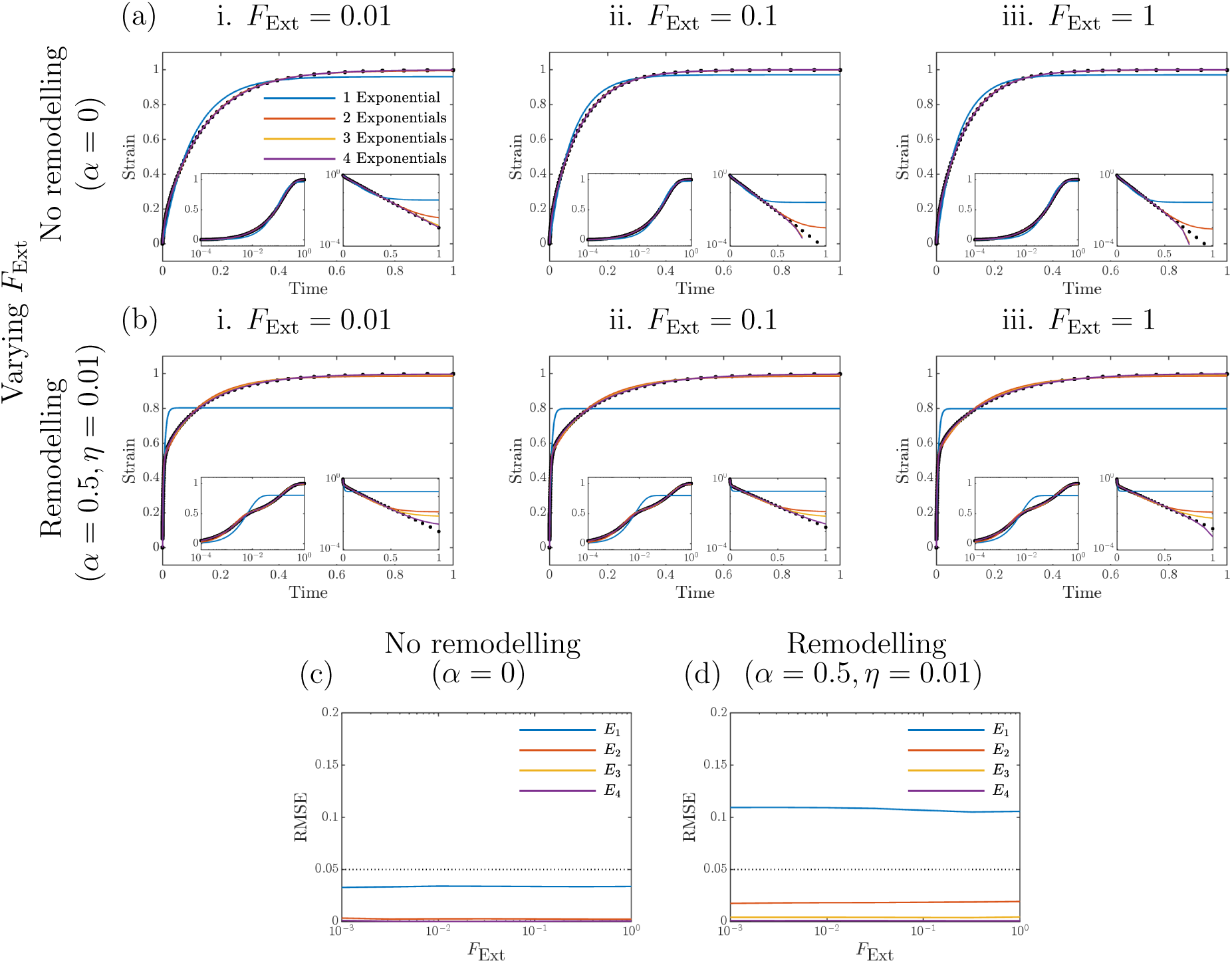
Applied force has no effect on the number of timescales in the creep experiment. Time–series for results presented in Figure 6 (a) and (b). (a) and (b) show the re–scaled strain over time of the tissue as the applied force *F*_Ext_ is varied, for no remodelling (*α* = 0) and with remodelling (*α* = 0.5 and *η* = 0.01), respectively. Simulation results are given by black dots, with exponential fits as shown in the legend. Insets show the strain with a log scale for time (left) and 1*−*strain with a log scale (right). (c) and (d) show the root mean square error of the exponential fits as *F*_Ext_ is varied without and with remodelling respectively. Dotted black lines in (c) and (d) show the error threshold of *E*_T_ = 0.05.

**Supplementary Figure S2:**
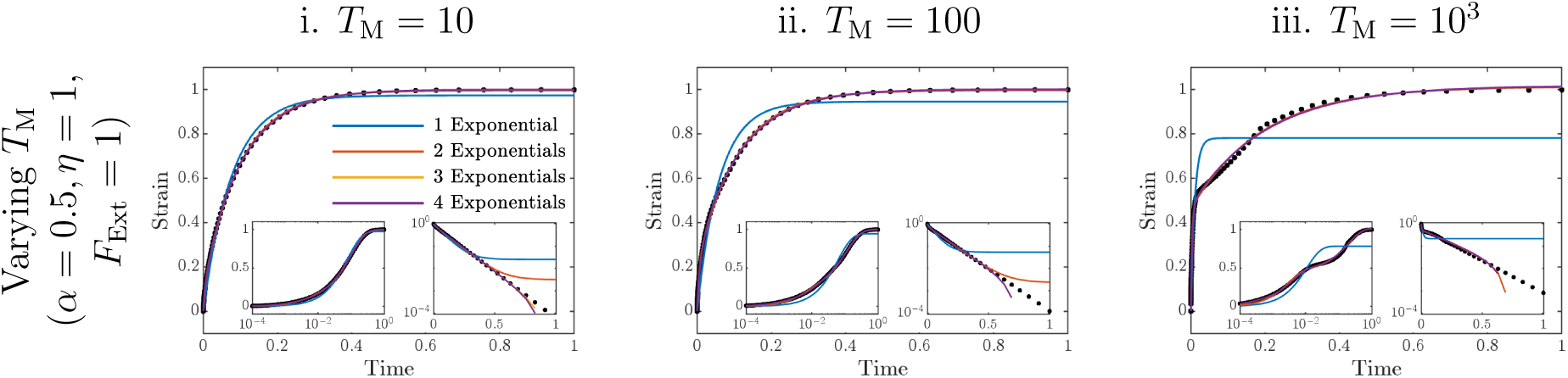
Memory leads to multiple timescales in the creep experiment. Time–series for results presented in Figures 6 (c)–(e). Re–scaled strain over time of the tissue under an applied force *F*_Ext_ = 1 for *α* = 0.5 and *η* = 1, with remodelling memories *T*_M_ = 10, 100, 10^3^. Simulation results are given as black dots, with exponential fits as shown in the legend. Insets show the strain with a log scale for time (left) and 1*−*strain with a log scale (right).

**Supplementary Figure S3:**
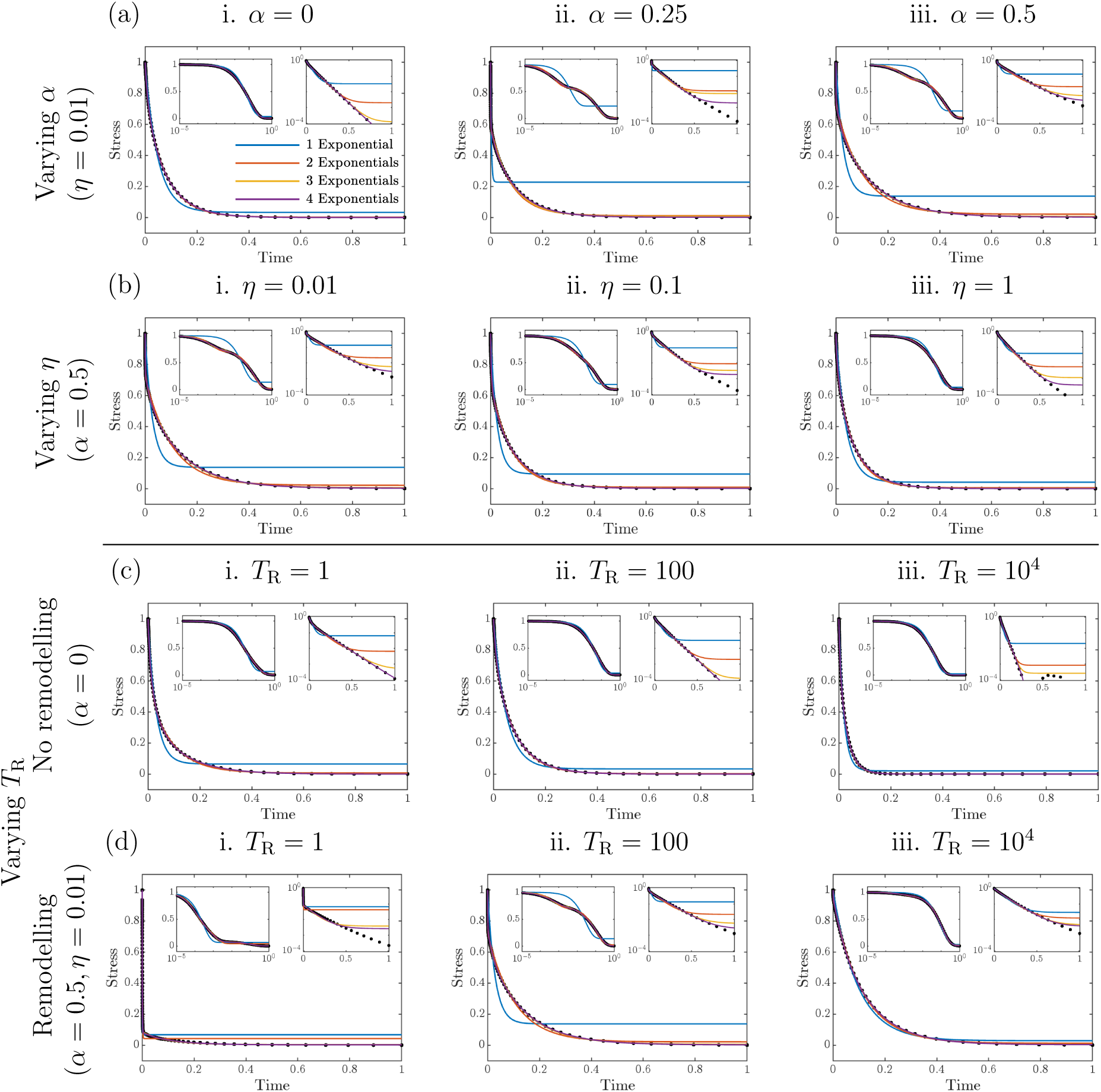
Varying dynamic parameters or the duration of applied strain can lead to multiple timescales in the stress relaxation experiment. Time–series for results presented in Figures 9 (a)–(d). (a) and (b) show the re–scaled stress of the tissue after an applied strain had been imposed over a time *T*_R_ = 100, as *α* and *η* are varied, respectively. (c) and (d) show the re–scaled stress of the tissue after an applied strain had been imposed over a time *T*_R_, as *T*_R_ is varied, for no remodelling (*α* = 0) and with remodelling (*α* = 0.5 and *η* = 0.01), respectively. Simulation results are given as black dots, with exponential fits as shown in the legend. Insets show the stress with a log scale for time (left) and with a log scale for stress (right). All simulations are with no memory, *T*_M_ = 0.

**Supplementary Figure S4:**
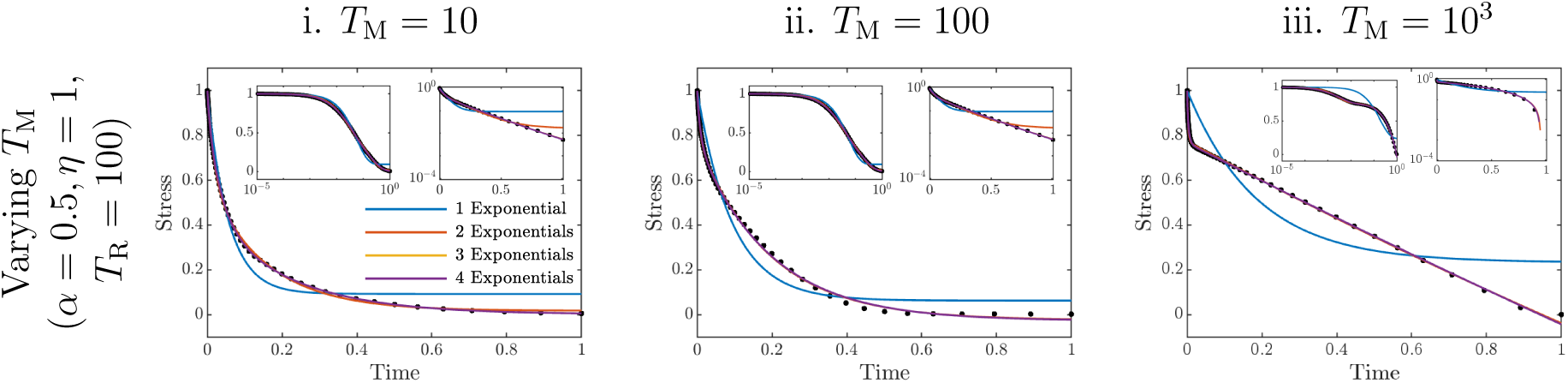
Memory leads to multiple timescales in the stress relaxation experiment. Time–series for results presented in Figures 10 (e) and (f). Re–scaled stress of the tissue after an applied strain had been imposed over a time *T*_R_ = 100, for *α* = 0.5 and *η* = 1, with remodelling memories *T*_M_ = 10, 100, 10^3^. Simulation results are given as black dots, with exponential fits as shown in the legend. Insets show the stress with a log scale for time (left) and with a log scale for stress (right).

## Footnotes

§ Note that the choice of 10*^−^*^6^ is somewhat arbitrary, but was chosen to be small enough to ensure that the system is close to equilibrium. We confirmed that varying this threshold by an order of magnitude in either direction does not significantly change the results presented here.

|| The choice *N* = 1001 was validated by confirming that increasing to *N* = 5001 produces a negligible change in the resulting fit errors, *E_n_*, reported here.

¶ An alternative threshold could be chosen based on the desired sensitivity of the classification, here we have used 0.05 to balance the trade–off between sensitivity and specificity. It was largely chosen as it identifies between the single and multiple timescales seen in Figure 4.

+ This is the same criterion as used for equilibrium in the creep relaxation experiments, but with the stress instead of the strain. Again if we change the threshold by an order of magnitude in either direction, the results presented here do not significantly change.

## Notes

### Competing Interest Statement

The authors have declared no competing interest.

